# Hierarchical organization of mechano-nociceptive pathways revealed by activity labeling

**DOI:** 10.64898/2025.12.10.693066

**Authors:** FM de-Faria, GB Carballo, MB Maidana Capitan, I Szczot, M Brodzki, C Karlsson, LF Nascimento, H Olausson, M Larsson, M Szczot

## Abstract

Noxious mechanical stimuli give rise to distinct percepts, from sharp cutaneous pain to diffuse visceral discomfort, yet the nociceptor ensembles that underlie these differences remain poorly defined. We mapped the peripheral architecture of nociceptive signaling by combining *in vivo* activity labeling of pelvic nerve afferents with single-cell RNA sequencing. Noxious stimuli recruited diverse classes of mechano-nociceptors, but stimulus type exerted modest influence on the composition of activated ensembles. Instead, tissue identity imposed the dominant organizational structure: stimulation of cutaneous and deep pelvic tissues engaged distinct subsets of both myelinated and unmyelinated neurons, revealing a clear domain-level division. Within this architecture, we identified a bladder-innervating myelinated nociceptor subtype with distinctive molecular features, illustrating an additional layer of refinement. Functional imaging and anatomical tracing corroborated this multilevel organization. These findings reveal a hierarchical organization of peripheral mechanical pain encoding, in which mechano-nociceptor populations are differentially engaged according to tissue domain and organ context.

## Introduction

Nociception safeguards organisms from external injury and internal tissue damage by detecting and distinguishing harmful stimuli, and encoding their intensity and location, thereby driving adaptive behavioral and homeostatic responses. While cutaneous noxious stimuli can elicit rapid withdrawal reflexes^1,2^, distension of visceral organs induces discomfort, defensive reflexes^3-5^ and urgency^5,6^. These diverse perceptual and physiological outcomes arise from the selective recruitment of ensembles of sensory neurons that reside in dorsal root ganglia (DRG) and innervate somatic and visceral tissues. Yet, the molecular organization of these ensembles across the somato–visceral axis and stimulus types *in vivo* remains poorly understood.

Organ-specific requirements for sensory feedback, together with differences in the local mechanical environment, create unique demands on nociceptive signaling across organ systems. These demands are particularly evident in lumbosacral DRGs, which give rise to pelvic nerve sensory afferents innervating urinary, digestive, reproductive, and somatic targets^7-11^. Moreover, mechanical stimuli themselves vary in intensity and type ranging from stroking to stretch, and activate distinct subsets of sensory neurons^3,12,13^. Delineating how these stimulus features are represented within the broader ensemble of responding neurons is essential to understanding the biological relevance of distinct neuronal classes and their interactions. Among neurons contributing to ensemble responses, well-studied cutaneous low-threshold mechanoreceptors (LTMRs) are specialized to encode gentle inputs^14-18^, whereas a distinct set comprises nociceptors that are unresponsive to gentle stimuli but respond robustly to high-intensity mechanical forces.

To understand the architecture of the nociceptive system, nociceptor populations have traditionally been categorized across multiple domains, including conduction velocity^19,20^, anatomical projection patterns^21-23^, neurochemistry^24^, and stimulus-response properties^1,12,25-27^. Over the past decade, molecular profiling through single-cell transcriptomics has transformed our understanding of sensory neuron organization, providing both a systems-level^28-30^ and an organ-specific view of cellular diversity^31^. Recently, integrated DRG atlases have extended this framework further, identifying additional molecular subtypes^32,33^.

The growing number of transcriptomically defined sensory neuron subtypes has raised fundamental questions about how stimuli are combinatorially encoded by nociceptor ensembles across the body. One strategy to address these questions is to employ multiple Cre driver lines to map the DRG transcriptomic space, enabling cell-type-specific identification of cutaneous^13,14^ or colonic^3^ afferents. A complementary approach has combined pan-neuronal calcium imaging with post hoc multiplexed fluorescent in situ hybridization (FISH) to link touch-evoked responses under physiological and inflammatory conditions to transcriptomic identity in trigeminal ganglia^34,35^. Although these approaches have yielded important insights, their reliance on predefined molecular markers biases sampling, limits the ability to resolve related nociceptor subtypes^30,32,33^ and does not allow system-level analysis of nociceptive coding across tissues and organs.

Therefore, to facilitate decoding of the cellular logic of nociceptive signaling, we developed an unbiased, activity-based strategy that integrates in vivo functional characterization using the photoconvertible calcium reporter CaMPARI^36^ with transcriptomic profiling. We applied this approach to map nociceptors activated by naturalistic noxious stimuli in the bladder, colorectum and caudal skin - organs innervated by the pelvic nerve. Post hoc single-cell RNA sequencing (scRNA-seq) of the labeled neurons revealed the molecular identities of the responsive populations, enabling us to map both known and novel nociceptor classes engaged during stimulation. Our findings reveal that nociceptive pathways are organized hierarchically, with molecular specificity increasing from fiber type to organ-specific subclasses.

## Results

### CaMPARI-based activity labeling reveals transcriptomic identities of stimulus-responsive neurons

Identifying nociceptor ensembles is challenging because activity-dependent genetic labeling tools, such as immediate-early gene reporters, are not reliably induced in DRG neurons. We therefore developed an optical approach using the genetically encoded Calcium Modulated Photoactivatable Ratiometric Integrator (CaMPARI^36^). CaMPARI’s green fluorescence is quenched by calcium influx in active cells; if this influx coincides with ultraviolet (UV) illumination, the fluorophore’s emission color changes permanently from green to red (Fig. 1A). We reasoned that *in vivo* UV conversion during stimulation would allow *ex vivo* selection of neurons representing the tested stimuli for transcriptomic profiling by single-cell RNA sequencing (scRNA-seq), enabling subsequent ensemble identification (Fig. 1B). To drive widespread CaMPARI expression in sensory neurons, we injected an adeno-associated viral vector encoding Cre recombinase neonatally into the conditional CaMPARI line (*R26*^*LSL-CaMPARI*^). This strategy, previously used to drive expression of other calcium indicators^34,35,37^, induced widespread CaMPARI expression in DRG neurons. To permit UV illumination *in vivo*, we employed a surgical approach for functional imaging in anesthetized mice^1,38^ to gain optical access to lumbosacral DRGs.

**Figure 1.**
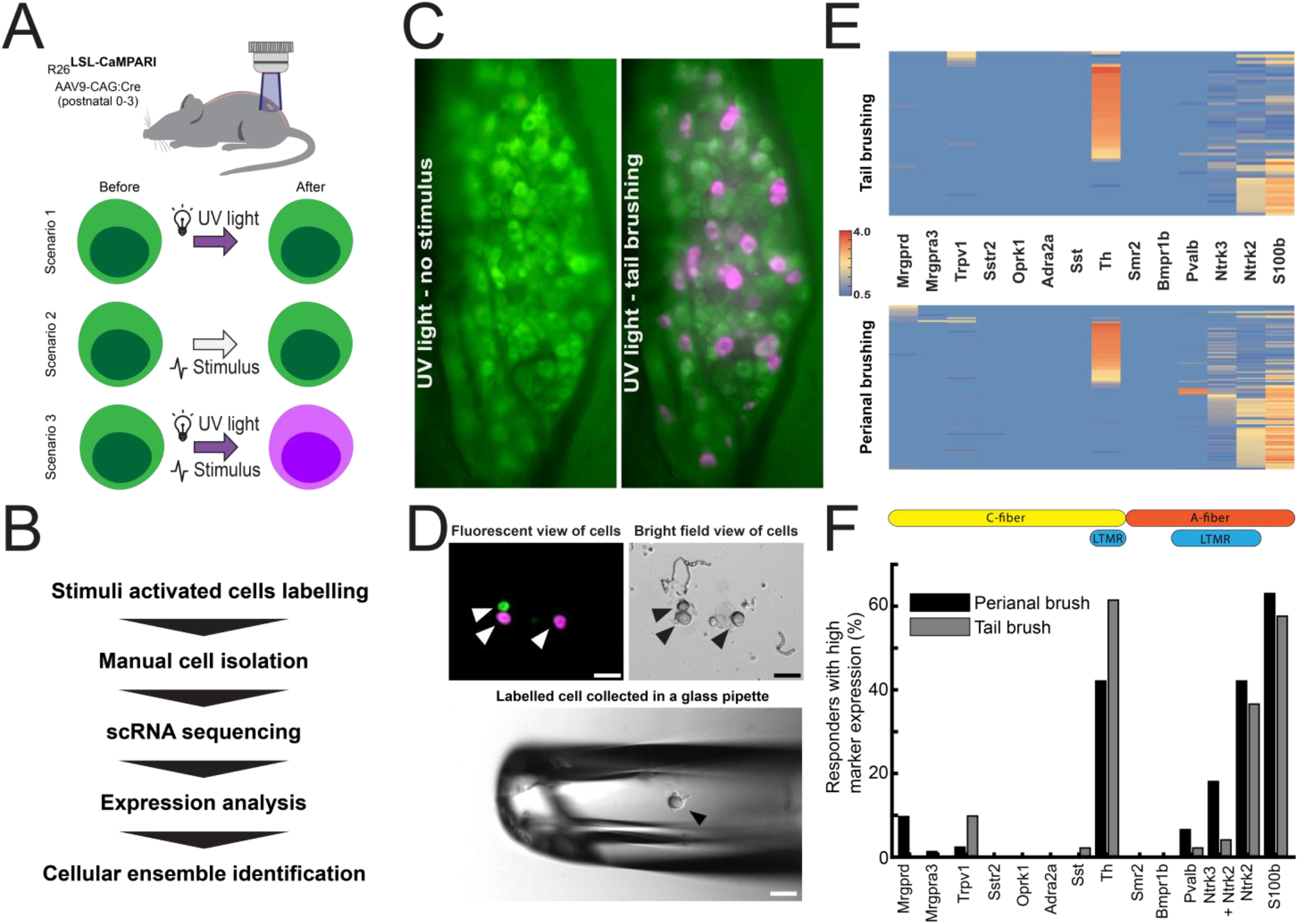
Framework for identifying neuronal ensembles activated by naturalistic stimuli. **A)** Schematic of the CaMPARI photoconversion strategy. Fluorophore conversion from green to red (pseudocolored magenta) occurs only when activity-induced Ca^2+^ elevation coincides with ultraviolet (UV) light exposure. **B)** Workflow for mapping stimuli-activated neuronal ensembles. **C)** Pelvic DRG in a CaMPARI-expressing mouse before (left) and after (right) photoconversion during tail brushing, showing basal green fluorescence and photoconverted cells (magenta) of the activated ensemble. **D)** Fluorescent and brightfield images of converted and non-converted cells, with a representative example of a collected cell. **E)** Heatmap of marker-gene expression in neurons responsive to perianal brushing (bottom; n = 89 cells, N = 6 mice) and tail brushing (top; n = 52 cells, N = 2 mice). Scale depicts log-normalized expression per cell. **F)** Expression of representative markers in brushing-responsive cells, indicating predominant activation of low-threshold mechanoreceptor (LTMR) populations, including unmyelinated (*Th*^*+*^) and myelinated (*Ntrk2*^+^ and *Ntrk2*^+^ *+ Ntrk3*^+^ double-positive) subtypes. Expression threshold was set to 1. *DRG: dorsal root ganglia; CaMPARI: calcium-modulated photoactivatable ratiometric integrator. Additional conversion characterization is provided in Fig. S1*.*1 and Fig. S1*.*2*.

As expected, we observed fluorescence dimming in a subset of DRG neurons during brushing of the perianal and tail skin, reporting neuronal activity (Fig. S1.1). Moreover, when stimulation was coupled with UV illumination, CaMPARI activity-dependent green-to-red photoconversion occurred in neurons that were active during live recording (Fig. S1.1). We further tested functional labeling of DRG neurons with CaMPARI across an array of naturalistic gentle and noxious stimuli and found it to be robust, reliable, and restricted to a subset of cells (Fig. 1C, Fig. S1.2). To control for photoconversion in spontaneously active cells, each experiment was preceded by a single UV illumination cycle without peripheral stimulation to assess baseline conversion. In most preparations, little to no photoconversion occurred under these mock conditions (Fig. S1.2). Preparations with detectable mock photoconversion (<5% of all preparations) were excluded from further analysis.

After establishing a conversion protocol, we proceeded to validate ensemble identification with activity labeling. We selected gentle brushing of hairy skin because its neuronal correlates have been extensively characterized, with existing evidence indicating predominant activation of C-LTMRs and Aδ-RA-LTMRs, along with a contribution from Aβ-RA-LTMRs^14,34,35^. We independently analyzed responses to gentle brushing at two distinct sites, the lateral base of the tail and the perianal region, predicting that both would engage similar distributions of sensory neuron subtypes. Both paradigms led to robust photoconversion (Fig. 1C, Fig. S1.1), and photoconverted cells were easily recovered for scRNA-seq under a fluorescent microscope (Fig. 1D), resulting in 52 and 89 cells in response to tail and perianal brushing, respectively.

Following scRNA-seq of activity-labeled neurons, we assessed the expression of molecular markers used to identify DRG neurons, based on established criteria^29,30,32,33^. As predicted for both types of cutaneous stroking, most labeled cells were identified as C-LTMRs (>40%, identified via *Th* expression), while majority of the remaining cells were myelinated Aδ-RA-LTMRs (>30%, identified based on co-expression of *S100b* and *Ntrk2* (Fig. 1E-F). The reproducible outcome demonstrates the robustness of our approach and reinforces the role of C-LTMRs in encoding gentle brushing. Additionally, a subset of myelinated neurons responding to perianal brushing co-expressed *Ntrk3* and *Ntrk2*, indicative of Aβ-RA-LTMRs^33^; their underrepresentation in tail brushing likely reflects regional innervation differences. Fewer than 10% of cells responding to gentle brushing expressed nociceptor markers *Mrgprd* (perianal brushing) or *Trpv1* (tail brushing). Notably, recent work^13^ has shown that some *Mrgprd*-positive neurons can exhibit mild brush sensitivity, especially in regions with dense pelvic innervation^39^. No cells expressing markers of other high-threshold mechanoreceptors (HTMRs) such as *Bmpr1b, Smr2, Mrgpra3, Sstr2, Oprk1*, or *Adra2a* were detected in this paradigm (Fig. 1E-F). Taken together, these findings demonstrate that our activity-labeling protocol reliably identifies neural ensembles.

### Pelvic sensory afferents exhibit unique functional and molecular organization

Correct interpretation of ensemble mapping requires detailed knowledge of the functional and transcriptomic context within pelvic nerve sensory afferents. First, we performed in-depth functional characterization of lumbosacral DRG neurons innervating pelvic targets using *in vivo* calcium imaging in a pan-neuronal GCaMP6f expression model generated by neonatal injection of an AAV carrying Cre into Ai95 (*R26*^*LSL-GCaMP6f*^) reporter mice^37^. We recorded calcium transients evoked by a panel of innocuous and noxious mechanical stimuli targeting bladder, colon, and skin (Fig. 2A, Fig S2.1).

**Figure 2.**
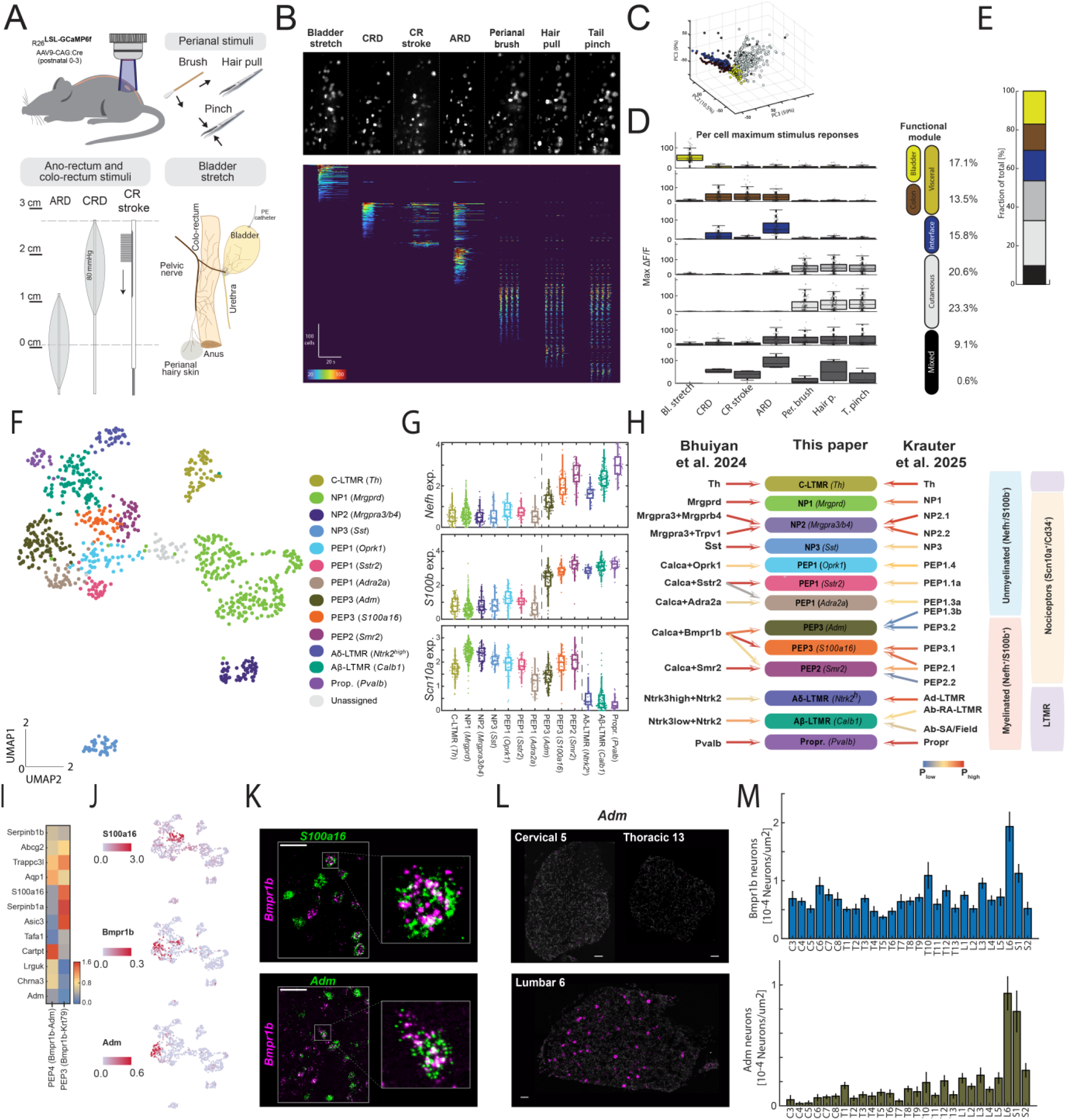
Functional and molecular characterization of pelvic sensory neurons. **A)** Experimental design for studying the functional organization of pelvic sensory neurons. GCaMP mice were imaged *in vivo* during application of a diverse set of naturalistic mechanical stimuli. Cutaneous stimuli included perianal brushing (Per. brush), hair pull (Hair p.), and base of tail pinch (T. pinch); gastrointestinal stimuli included anorectal distension (ARD), colorectal distension (CRD), and colorectal stroking (CR stroke); urinary system was probed through bladder stretch (Bl. stretch). **B)** The top panel shows a standard-deviation projection in grayscale highlighting stimulus-responsive cells in a representative experiment (see also Fig. S2.1). The bottom panel presents a heatmap summarizing all recorded responses (n = 666 neurons, N = 5 mice) across the full set of stimuli. Cells are ordered by their preferred stimulus to visualize functional tuning. **C)** Principal component distribution of individual cells’ response vectors. Response vectors were constructed from maximum responses to each stimulus. Color-coding reflects cells’ functional module assignment obtained from representative Gaussian mixture model (GMM) clustering. **D)** Most representative GMM clustering of pelvic sensory neuron response vectors revealed anatomically segregated modules, with cells in each module showing distinct target specificity. Bars represent maximal single-cell responses to each stimulus for all neurons classified within the corresponding cluster. A distinct cluster was identified that selectively mediates responses to the stimuli at the cutaneous–visceral interface (ARD), whereas few neurons were classified into clusters with mixed responses (see also Fig. S2.2) **E)** Stacked bar graph quantifying the distribution of neurons across functional modules identified in **D. F)** Uniform Manifold Approximation and Projection (UMAP) of control pelvic sensory neurons profiled by Smart-Seq3 scRNA-seq. Colors indicate clusters identified by the Louvain algorithm, and annotated based on gene expression analysis (see also Fig. S2.3-S2.5). **G)** Differential expression analysis of myelination (*Nefh, S100b*) and nociceptor (*Scn10a*) markers across the clusters identified in **F**. Scale depicts log-normalized expression per cell. **H)** Summary of relationships between the classes identified in this study and those reported in the largest available consensus atlases (Bhuiyan et al., 2024; Krauter et al., 2025; see also Fig. S2.6). Relationships were derived by embedding control cells into the consensus atlases. Arrow colors indicate posterior embedding probability and assignment confidence. **I)** Expression matrix of marker genes shared by, and distinguishing between, the newly identified PEP3 subclasses. Color scale indicates normalized log-transformed expression levels. **J)** UMAP highlighting the expression of markers identifying PEP3 classes. **K)** Fluorescent in situ hybridization (FISH) analysis showing expression of *Adm* and *S100a16* within *Bmpr1b*^+^ neurons. Images demonstrate *Adm*^+^ and *S100a16*^+^ subsets of Bmpr1b^+^ neurons. Scale bar 50μm. **L-M)** FISH analysis of Bmpr1b^+^ and Adm^+^ neuron abundance along the neuroaxis. Bmpr1b^+^ neurons expressed uniformly across all levels except lower lumbar. In contrast, Adm^+^ neurons are highly enriched in L6 and S1 ganglia, with sparse representation at more rostral levels (n≥6, scale bar 50μm, see also Fig. S2.7).

To probe the urinary system, a catheter was inserted into the bladder dome, allowing controlled saline perfusion to stretch the bladder. For colorectal stimulation, a barostat-controlled mini-balloon delivered noxious colorectal distension (CRD, 80 mmHg), followed by a rostro-caudal stroke along the colorectum using a customized micro-brush (CR stroke). Anorectal distension (ARD) was induced by positioning the mini-balloon halfway into the anal sphincter to engage the cutaneous–visceral interface. Finally, to probe cutaneous inputs, we used gentle brushing to engage LTMRs, followed by hair-pull and a pinch at the hairy skin–covered base of the tail to probe nociceptive responses.

After recording the responses to all stimuli, we first analyzed the stimulus selectivity for each cell (Fig. 2B, Fig. S2.1) and then, for every activated cell, we quantified the maximum calcium transient for each stimulus and constructed multidimensional response vectors. We subjected these vectors to principal component analysis (Fig. 2C) and repeatedly applied unsupervised Gaussian Mixture Modeling (50 000 repetitions, Fig. S2.2) to find the most representative GMM cluster assignment and identify functionally related clusters of cells. This analysis showed that neurons clustered into functional modules tuned to specific tissue domains: visceral neurons segregated into colon- and bladder-specific modules, whereas cutaneous stimuli typically recruited only skin-responding cells (Fig. 2D-E). Anorectal distension was an exception, as it activated neurons within a dedicated module as well as subsets within visceral and cutaneous modules, consistent with the mixed sensory anatomy at the cutaneous–visceral interface. Thus, overall, pelvic sensory neurons are organized into functionally distinct modules, with bladder and colon inputs showing clear segregation and limited overlap with cutaneous inputs. This architecture is most evident under high-intensity stimulation, when dedicated visceral and somatic pathways are strongly recruited.

Next, we extended our characterization to the transcriptomic landscape. To this end, we manually collected ∼1000 cells for scRNA-seq to create a control atlas (Fig. 2F). Using the SmartSeq3 pipeline^40^, we achieved deeper coverage than typical available atlases, with ∼100,000 unique molecular identifiers (UMIs) and ∼10,000 genes detected per cell, and quality control analyses demonstrated high fidelity of the atlas as well as robustness of unbiased clustering (Fig. S2.3). Following marker analysis, we identified 13 clusters characterized by specific gene expression (Fig. 2F, Fig. S2.3C-F, Fig. S2.4, Fig. S2.5). Most clusters could be correlated with described neuronal classes based on established markers, allowing us to apply standardized nomenclature^29^. To capture additional diversity within broader classes, we also included the characteristic marker in each annotation label. Based on the expression of established markers (*Nefh, S100b, Thy1*), we categorized cells into myelinated and unmyelinated groups and identified nociceptors (*Scn10a*^*+*^ *Cd34*^*-*^)^29,32^ (Fig. 2G,H). Our atlas did not capture some smaller populations such as *Trpm8*^*high*^ thermoreceptors^28^ or *Dcn*^*+*^ cells^33^, possibly reflecting their scarcity in lumbosacral ganglia or the resolution limits of our atlas. While most of the neuronal classes had counterparts in previously published DRG atlases, others appeared to represent new subdivisions (e.g. PEP3 classes), and others corresponded to neuron types recognized as molecularly diverse but without well-defined functional specialization (e.g. PEP1). To better resolve cellular identities, we compared our clusters with those in recent harmonized DRG atlases integrating multiple datasets. We integrated our data with the consensus dataset from Bhuiyan et al.^33^ and Krauter et al.^32^ and calculated the average probability of classifying cells from our dataset in their clusters (Fig. 2G, Fig. S2.6); most populations mapped to their predicted counterparts. However, we noticed a split within the PEP3 population into PEP3 (*S100a16*) and PEP3 (*Adm*), a subdivision previously reported solely in the high-fidelity atlas^32^ (Fig. 2H). Integration with the Krauter et al.^32^ dataset further suggested that the PEP3 (*Adm*) group may harbor additional subdivisions, though these were not clearly delineated in unbiased clustering of our atlas. The observation of two separate clusters within the *Bmpr1b*^*+*^ population of PEP3 nociceptors was supported by expression analysis showing divergent marker profiles (Fig. 2I-J) and by FISH (Fig. 2K). We hypothesized that PEP3 (*Adm*) may represent a pelvic-enriched population not well represented in previous datasets^30,33^ derived from neurons pooled across multiple spinal levels. Marker expression along the neuraxis confirmed enrichment of *Adm*^*+*^ neurons in L6–S1 ganglia, coinciding with an increased density of *Bmpr1b*^*+*^ neurons in the same region (Fig. 2L-M, Fig. S2.7). We also compared this with C-fiber PEP1 (*Adra2a*) neurons, which are known to be depleted in ganglia with increased representation of cutaneous tissues^13^. Interestingly, in contrast to PEP1 (*Adra2a*), which is found across multiple spinal levels, PEP3 (*Adm*) was very sparsely expressed beyond L6–S1 ganglia (Fig. S2.7). Together, these data reveal functional ensembles and organ-specific pelvic-enriched populations, highlighting distinct sensory specialization within lumbosacral ganglia.

### Mechanical stimuli recruit multiple classes of sensory neurons

To analyze ensemble coding across innocuous and noxious stimulus modalities, we embedded stimulus-labeled cells into the atlas and broadly categorized them into four classes defined by myelination and nociceptor markers (Fig. 2G-H, Fig. 3A). For cross-modality characterization, we compared representative gentle stimuli we mapped initially (perianal and tail brushing, Fig. 1) with noxious cutaneous (tail pinch) and visceral stimuli (colorectal stroke, Fig. 3B-C). Cells representing each stimulus ensemble were collected from multiple preparations (Supplementary Table 1) and subjected to the same quality check as control cells. Gentle stimuli were predominantly encoded by LTMRs, with A-LTMRs comprising 30–40% and C-LTMRs 30–50% of activated neurons (Fig. 3D, Supplementary Tables 1-3). The remainder reflected minor activation of nociceptive C-fibers, consistent with limited brush sensitivity reported in some of these cells^1,13^. Next, we assessed encoding of noxious stimuli across tissues using pinch (cutaneous) and colorectal stroke (visceral) as examples. Both activated substantial A-LTMR populations, but visceral stimulation elicited minimal C-LTMR responses. As expected, noxious stimulation increased nociceptor recruitment, with A-fiber nociceptors emerging only under noxious conditions and nociceptors overall comprising 50– 80% of activated C-fibers and ∼40% of activated A-fibers (Fig. 3D–E). These data demonstrate that even though noxious stimulation activates LTMRs, the shift in stimulus valence is driven by the co-recruitment of nociceptors into the ensemble. In particular, transitioning from gentle to noxious stimuli is accompanied by progressive recruitment of C-fiber nociceptors and the appearance of fast-conducting A-fiber nociceptor activity.

**Figure 3.**
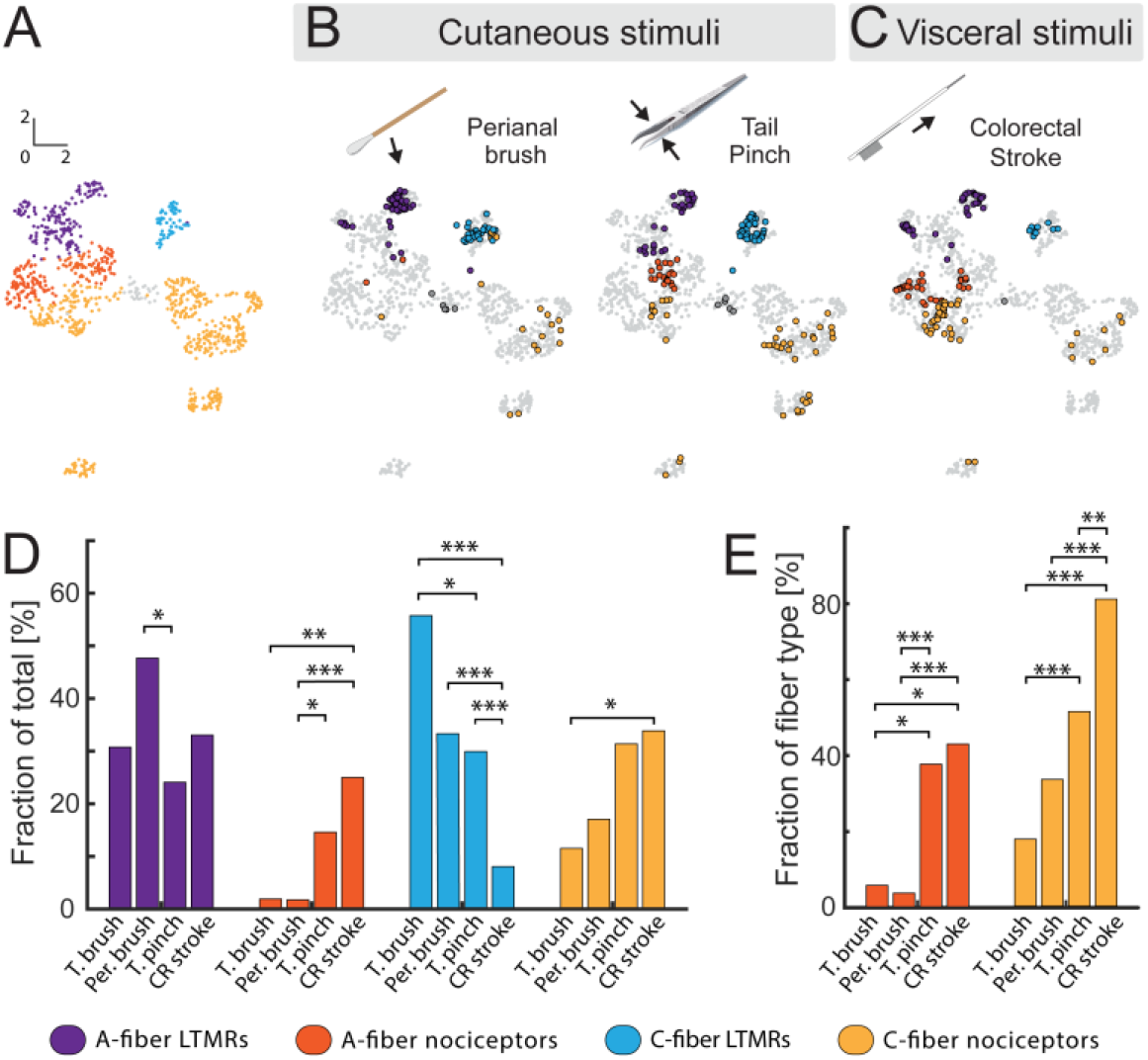
Activity labeling reveals organization of encoding cutaneous and visceral mechanical stimuli. **A)** Uniform Manifold Approximation and Projection (UMAP) of pelvic sensory neuron atlas highlighting overarching neuronal classes defined by canonical markers and pooled Louvain clustering. Classes include A-fiber low-threshold mechanoreceptors (LTMRs, *Nefh*^*+*^, *Scn10a*^*-*^), A-fiber nociceptors (*Nefh*^*+*^, *Scn10a*^*+*^), C-fiber LTMRs (*Nefh*^*-*^, *Scn10a*^*+*^, *Cd34*^*+*^), C-fiber nociceptors (*Nefh*^*-*^, *Scn10a*^*+*^, *Cd34*^*-*^). **B), C)** Overlay of the pelvic DRG transcriptomic atlas (gray) with labeled stimulus-responsive neurons, color-coded by their class membership. **B)** Cells activated by cutaneous gentle (perianal brush) and noxious stimuli (tail pinch). **C)** Cells activated by visceral noxious stimulation (colorectal stroke). **D)** Bar plots showing ensemble composition for gentle and noxious stimuli, demonstrating preferential engagement of C-fiber and A-fiber nociceptors during noxious cutaneous and visceral stimulation. **E)** Bar plots showing recruitment of nociceptors among A-fiber and C-fiber populations during cutaneous and visceral stimulation. *\*p < 0*.*05, **p < 0*.*01, ***p < 0*.*001; likelihood-ratio test with post hoc Fisher’s exact test (Holm correction). Sample sizes and tests statistics in Supplementary Tables 1–3*.

### Cutaneous and visceral noxious stimuli recruit different unmyelinated nociceptor ensembles

Unmyelinated C-nociceptors are important for affective aspects of pain^41,42^ as well as for autonomic functions along the gut–brain axis^4,43^. To probe the organ- and stimulus-specific organization of C-nociceptor ensembles, we expanded our analysis to include two additional noxious stimuli, hair-pulling and anorectal distension, thereby covering a range of stimuli that recruit visceral, cutaneous, and interface ensembles (Fig. 2D-E, Fig. S4.1). Cutaneous stimuli (pinch and hair-pull) activated a similar ensemble of C-nociceptors. Most responsive cells belonged to non-peptidergic classes, with 50– 70% identified as NP1 (*Mrgprd*) and a substantial fraction as NP2 (*Mrgpra3/b4*), so that non-peptidergic neurons accounted for 70–80% of the C-nociceptors (Fig. 4B-C, Fig. S4.2, Supplementary Tables 4-6). Among the remaining peptidergic neurons, the most prominent activation was observed in the PEP1 (*Oprk1*) class, although their overall contribution to the ensemble remained limited (Fig. 4C-D). The activation profile for anorectal distension was similar to that observed for the cutaneous stimuli, possibly reflecting dense C-fiber innervation in the perianal region^39^. However, colorectal stroking responses were dominated by a combination of peptidergic classes, with over 60% of the activated ensemble comprising PEP1 (*Adra2a*) and PEP1 (*Oprk1*) cells.

**Figure 4.**
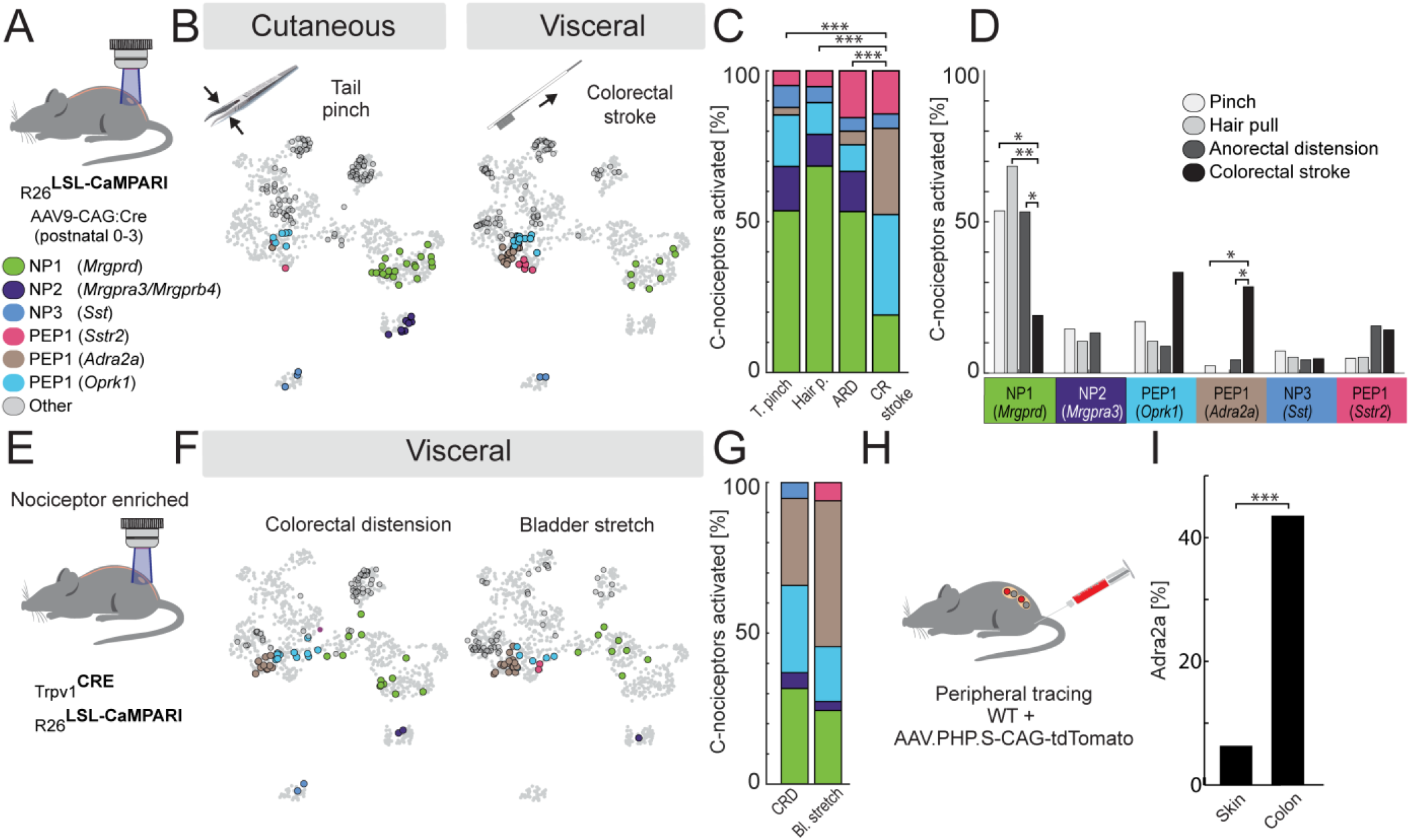
C-nociceptors differentially encode visceral and cutaneous noxious stimuli. **A)** Experimental strategy using the virally induced CaMPARI mouse model. Representative populations of DRG neurons were labeled by perinatal injection of an AAV vector encoding Cre recombinase, inducing CaMPARI expression in infected cells. **B)** Overlay of the pelvic DRG transcriptomic atlas (gray) with labeled stimulus-responsive C-nociceptors, color-coded by transcriptomic class. **C) D)** Bar plots showing C-nociceptors ensemble composition for noxious stimuli encoding, demonstrating preferential engagement of NP1 (Mrgprd) cells for cutaneous and PEP1 (*Adra2a*) and PEP1 (*Oprk1*) for the visceral stimuli. *Hair p. – Hair pulling, ARD – Anorectal distension, CR stroke – Colorectal stroke* **E)** Experimental strategy to generate nociceptor-enriched reporter model, restricting CaMPARI expression to Trpv1-lineage neurons. **F)** Overlay of the pelvic DRG transcriptomic atlas (gray) with labeled stimulus-responsive C-nociceptors, color-coded by their class membership. **G)** Bar plots showing C-nociceptors ensemble composition for noxious stimuli encoding confirming the critical role of PEP1 (*Adra2a*) in visceral C-nociceptors ensemble. *CRD – Colorectal distension, Bl. Stretch – bladder stretch*. **H)** Experimental strategy for peripheral tracing of target-specific innervation. Viral vector was peripherally injected into the colon or skin to label DRG neurons projecting to these tissues. **I)** PEP1 (*Adra2a*) neurons are enriched in the colon. Bar graph quantifies the proportion of peripherally traced C-nociceptor neurons from the skin (2 of 32) and colon (5 of 13) that were classified as PEP1 (*Adra2a*) by scRNA-seq. *\*p < 0*.*05, **p < 0*.*01, ***p < 0*.*001; likelihood-ratio test with post hoc Fisher’s exact test (Holm correction). Sample sizes and test statistics are provided in Supplementary Tables 4-6*.

To investigate whether this pattern generalizes across viscera, we generated *Trpv1*^*Cre*^*;R26*^*LSL-CaMPARI*^ mice to enrich CaMPARI expression in nociceptors, as defined by developmental Trpv1 expression^1,34^. Using this model, we mapped responses to bladder and colorectal distension (Fig. 2A, Fig. 4E-F); both produced activation patterns similar to colorectal stroke, with ensembles dominated by recruitment of PEP1 (*Adra2a*) neurons and co-activation of PEP1 (*Oprk1*) cells (Fig. 4G). Finally, we sought to confirm whether, as previously suggested^13^, PEP1 (*Adra2a*) neurons specifically project to visceral organs. To test this, we injected AAV-PHP.S-tdTomato^44^ into either skin or colorectum and sequenced the labeled cells (Fig. 4H, Fig. S4.3). Indeed, PEP1 (*Adra2a*) cells were abundant among colon-innervating neurons but nearly absent among skin-innervating populations (Fig. 4I). Our results demonstrate a qualitative difference in C-nociceptor encoding between cutaneous and visceral tissues and reveal a generalizable role for PEP1 (*Adra2a*) neurons in visceral signaling. functional organization of afferents through calcium

### Cutaneous and visceral noxious stimuli recruit different myelinated nociceptor ensembles

Myelinated nociceptors innervate both cutaneous and visceral tissues^19,45^ and mediate a range of avoidance responses^2,3,5,13,46^, but little is known about their molecular organization and specialization across tissues. Therefore, to complete our dissection of the nociceptive system, we analyzed the ensemble coding of myelinated nociceptors. Again, we began with identifying imaging; to this end we generated a mouse line with expression of GCaMP6f restricted to myelinated afferents (*Nefh*^*CreERT2*^*;R26*^*LSL-GCaMP6f*^, Fig. 5A^38^) and characterized neuronal responses to the same panel of noxious stimuli as previously (Fig. 5B, Fig. S5.1). Unbiased clustering of activated neurons (Fig. 5C-D, Fig. S5.2) revealed groupings into functional modules like those in the general population (Fig. 2). Bladder- and colon-responsive neurons segregated clearly, whereas myelinated neurons responding to anorectal distension did not form a distinct emergent cluster (Fig. 5C-D).

**Figure 5.**
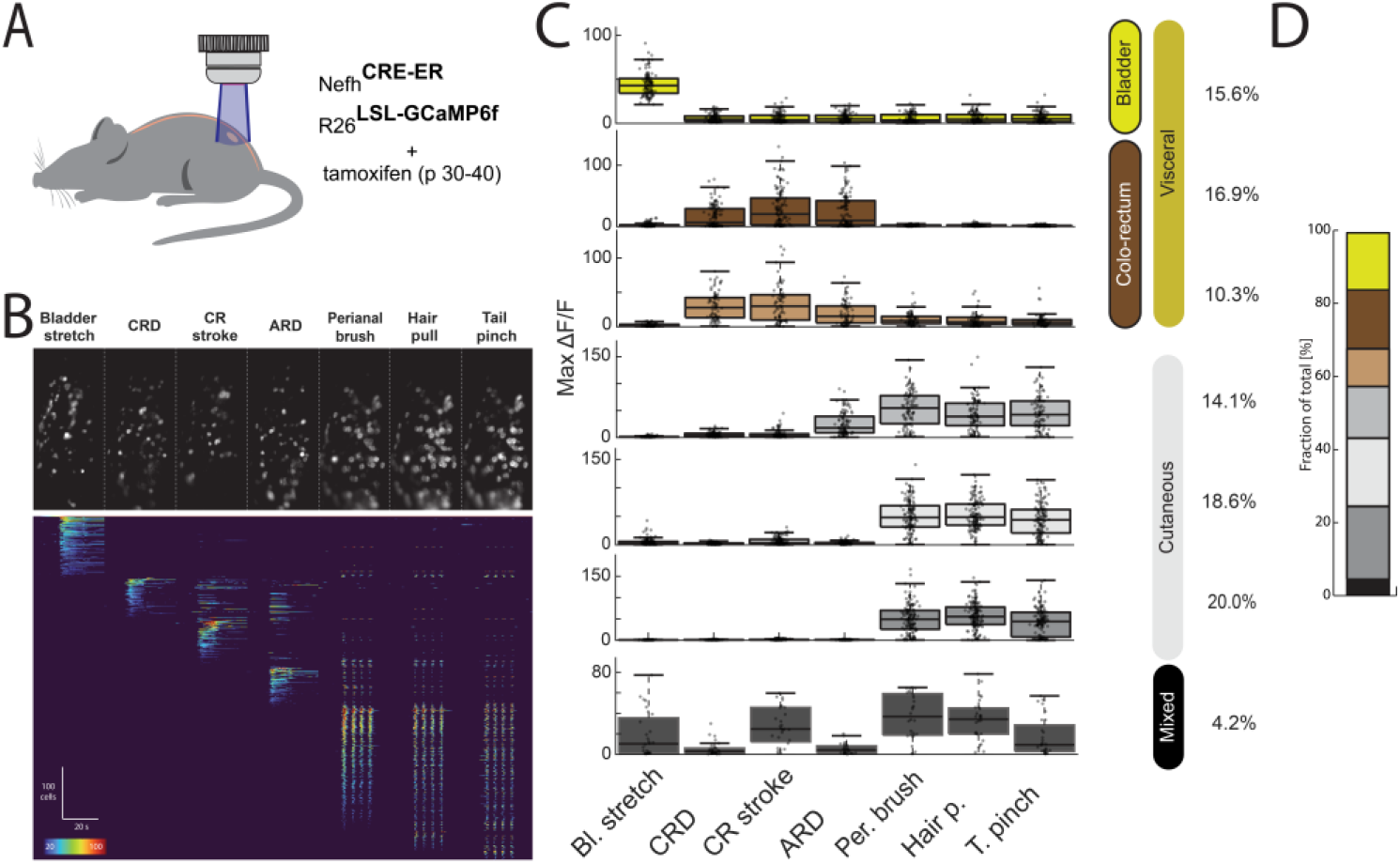
Functional characterization of myelinated pelvic sensory neurons. **A)** Experimental design for studying the functional organization of myelinated pelvic sensory neurons. A-fiber selective Nefh^CreER^; R26^LSL-GCaMP^ mice were imaged *in vivo* during application of a diverse set of naturalistic mechanical stimuli. Cutaneous stimuli included perianal brushing (Per. brush), hair pull (Hair p.), and pinch (T. pinch); gastrointestinal stimuli included anorectal distension (ARD), colorectal distension (CRD), and colorectal stroking (CR stroke); urinary system was probed through bladder stretch (Bl. Stretch). (see Fig. 2 for details). **B)** The top panel shows a standard-deviation projection in grayscale highlighting stimulus-responsive cells in a representative imaging experiment (see also Fig. S5.1). The bottom panel presents a heatmap summarizing all recorded responses (n = 665 neurons, N = 5 mice) across the full set of stimuli. Cells are ordered by their preferred stimulus to visualize functional tuning. **C)** Representative Gaussian mixture model (GMM) clustering of myelinated pelvic sensory neuron response vectors revealed anatomically segregated modules, with cells in each module showing distinct target specificity. Bars represent maximal single-cell responses to each stimulus for all neurons classified within the corresponding cluster. Note the absence of a cluster selectively grouping cutaneous– visceral interface (ARD)-responding cells, and the high proportion of cells within cutaneous modules. See also Fig. S5.2. **D)** Stacked bar graph quantifying the distribution of neurons across functional modules detected in **C**.

Next, we sought to quantify the class distribution of A-nociceptors activated in response to the stimuli (Fig. 6A-B, Supplementary Tables 4, 7-8). Of the three classes identified in our atlas (PEP3 (*S100a16*), PEP3 (*Adm*), and PEP2 (*Smr2*)), only PEP2 (*Smr2*) cells were not strongly activated by any stimuli. Strikingly, the remaining A-nociceptors showed clear target-specific specialization: PEP3 (*S100a16*) cells were tuned to cutaneous stimuli, while PEP3 (*Adm*) preferentially responded to colorectal stroke and anorectal distension (Fig. 6C-D). This distinction was further supported by data from the nociceptor-enriched model, where colorectal distension and bladder stretch mainly activated PEP3 (*Adm*) neurons (Fig. 6E–G), confirming their identity as specialized visceroceptors. myelinated Next, we examined whether this functional dichotomy mirrors anatomical organization. However, we reasoned that detection of peripherally traced PEP3 cells in DRG sections through FISH would be challenging. Therefore, we developed an alternative strategy where colorectum and the perianal skin were injected with Alexa Fluor-555 conjugated cholera toxin (CTB-555) in an intersectional transgenic line that labels myelinated nociceptors with mCitrine (*Nefh*^*CreER*^*;Scn10a*^*FlpO*^*;R26*^*LSL-FSF-ReachR-mCitrine* 38,47^). Subsequently, we dissociated the DRG and collected co-labeled cells for scRNA-seq. In line with our prediction, the general marker of PEP3, *Bmpr1b*, was present at a similar level in A-nociceptors innervating the skin and colon, while *Adm* expression was highly enriched in the colorectum-projecting population (Fig. 6H-I). Thus, A-nociceptors segregate into defined populations according to their target domains (cutaneous versus visceral).

**Figure 6.**
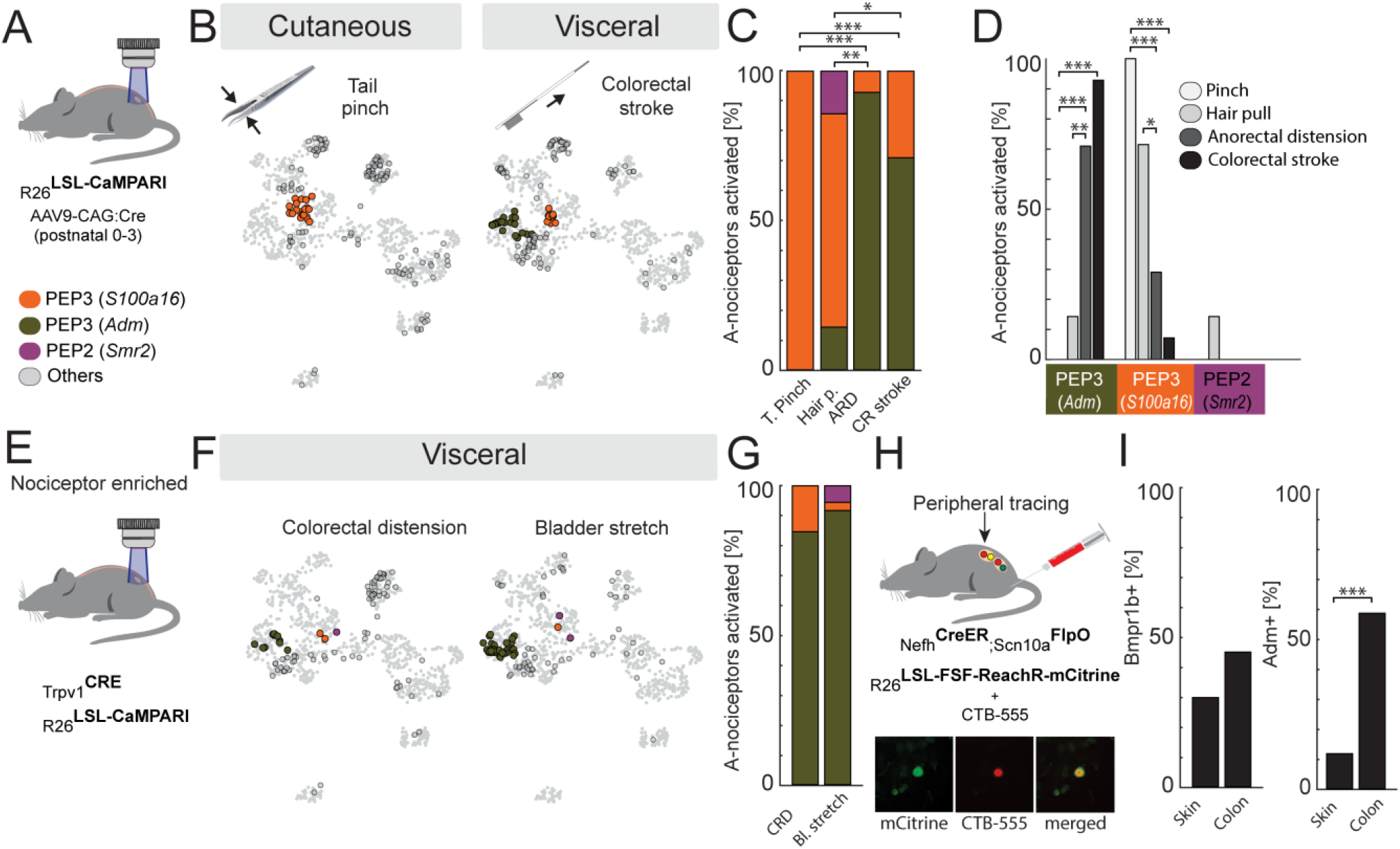
A-nociceptors differentially encode visceral and cutaneous noxious stimuli. **A)** Experimental strategy using the CaMPARI mouse model, as shown in Fig. 4A. The experiments were re-analyzed to illustrate A-nociceptor responses. **B)** Overlay of the pelvic DRG transcriptomic atlas (gray) with labeled stimulus-responsive A-nociceptors, color-coded by their class membership in the CaMPARI model. **C) D)** Bar plots showing A-nociceptors ensemble composition for noxious stimuli encoding, demonstrating preferential engagement of PEP3 (*S100a16*) cells for cutaneous and PEP3 (*Adm*) for the visceral stimuli. **E)** Experimental strategy to generate nociceptor-enriched reporter model, restricting CaMPARI expression to Trpv1-lineage neurons, as shown in Fig. 4E. The experiments were re-analyzed to illustrate A-nociceptor responses. **F)** Overlay of the pelvic DRG transcriptomic atlas (gray) with labeled stimulus-responsive A-nociceptors, color-coded by their class membership. **G)** Bar plots showing A-nociceptors ensemble composition for noxious stimuli encoding confirming the critical role of PEP3 (*Adm*) in visceral A-nociceptors ensemble. **H)** Experimental strategy for peripheral tracing of A-nociceptor innervation. CTB-Alexa Fluor 555 was peripherally injected into the colon or skin of *Nefh*^*CreER*^*;Scn10a*^*FlpO*^*;R26*^*LSL-FSF-ChR-mCitrine*^ mice to label A-nociceptor neurons projecting to these tissues. Double-labeled neurons (mCitrine^+^/CTB^+^) were collected and sequenced after colon or skin injections (bottom). **I)** While *Bmpr1b* expression was detected at similar levels in both skin- and colon-innervating A-nociceptor populations, *Adm* expression was largely restricted to visceral-innervating A-nociceptors (p = 3.42 × 10^−4^, Fisher’s exact test). *\*p < 0*.*05, **p < 0*.*01, ***p < 0*.*001; likelihood-ratio test with post hoc Fisher’s exact test (Holm correction). Sample sizes and test statistics in Supplementary Tables 4, 7, 8*.

### Myelinated nociceptors show organ specific-tuning

It has been proposed that sensory neuron identity reflects both developmental programs and peripheral cues^30^. We therefore asked whether molecular tuning in the somatosensory system could extend beyond broadly defined body domains to establish organ-specific signatures. To test this, we analyzed gene expression in visceral PEP1 (*Adra2a*) and PEP3 (*Adm*) neurons activated by bladder stretch or colorectal distension. PEP1 (*Adra2a*) neurons did not differ between bladder- and colon-responsive groups, whereas among PEP3 (*Adm*) we identified a distinct Uts2b^+^ subpopulation activated exclusively by bladder stretch (Fig. 7A). If *Uts2b*^*+*^ neurons preferentially innervate the bladder, they should localize to bladder-innervating ganglia. Indeed, FISH across sensory ganglia along the neuraxis revealed an accumulation of *Uts2b*^+^ neurons in the lower lumbar and sacral levels, with a strong peak at L6 (Fig. 7B). To confirm that PEP3 (*Uts2b*) neurons represent a distinct subset of the parent PEP3 class that projects specifically to the bladder, we used *Uts2b*^*Cre* 53^ and *Bmpr1b*^*Cre* 36^ intrathecally injected with AAV-CAG-Flex-tdTomato to investigate peripheral anatomy of these nociceptors (Fig. 7C). When we analyzed peripheral labeling, Bmpr1b^tdTomato^ fibers were detected in perianal skin, bladder, and colorectal tissues, whereas Uts2b^tdTomato^ fibers were confined to the bladder (Fig. 7C). Taken together, these data show that within a shared broad body domain, sensory neurons undergo further genetic specialization to support organ-specific encoding.

**Figure 7.**
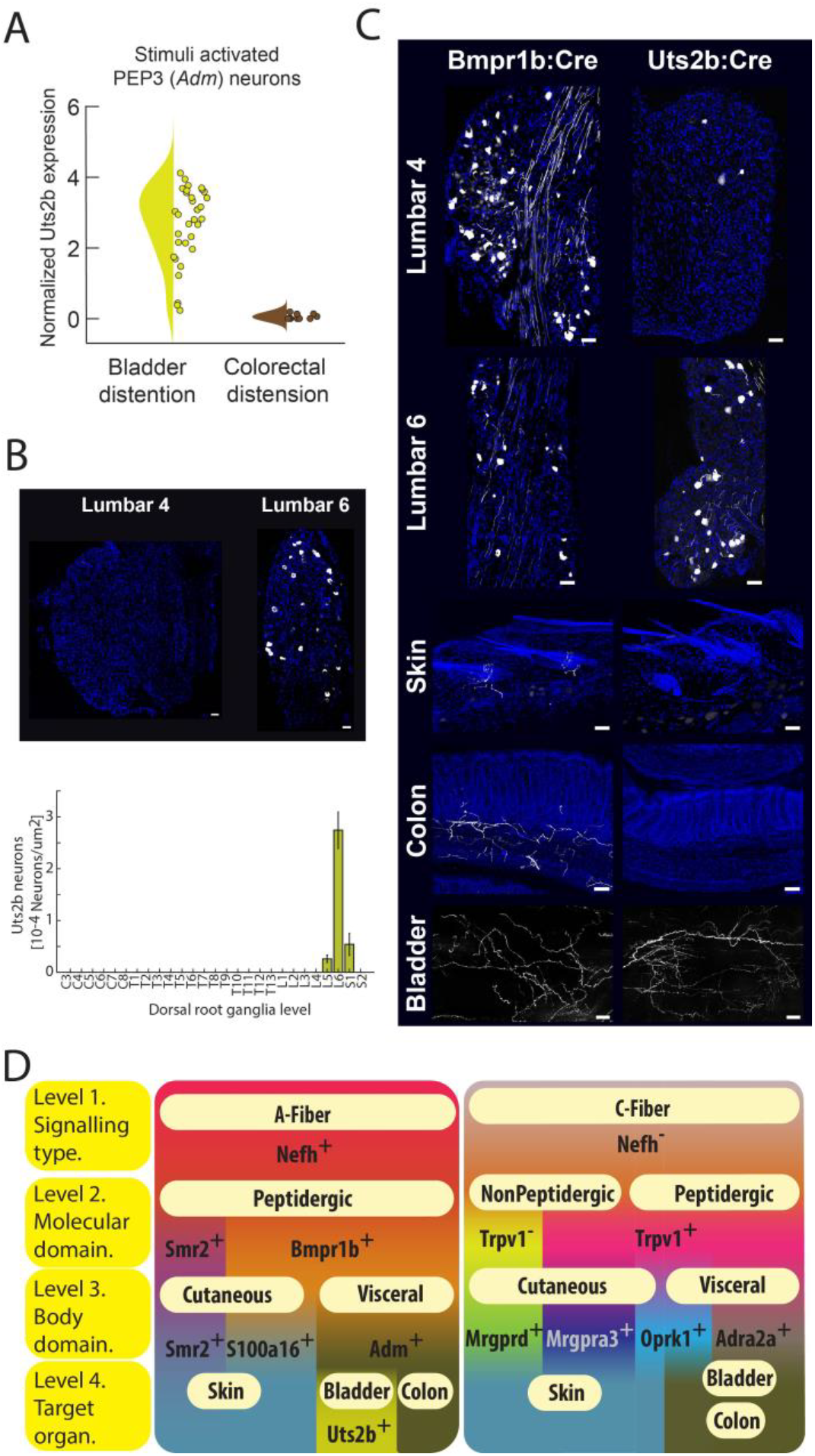
Uts2b defines bladder-specific innervation by A-nociceptors. **A)** PEP3 (Adm) neurons activated by bladder stretch express high levels of *Uts2b*. **B)** Fluorescence in situ hybridization analysis of *Uts2b*^+^ neurons abundance along the neuroaxis. Uts2b^high^ neurons are restricted to lumbar six ganglia, with sparse representation elsewhere (n≥6, 50μm). **C)** Uts2b-tdTomato labeling was limited to the bladder, whereas Bmpr1b-tdTomato labeling as expected showed projections to the colorectum, bladder, and perianal skin. **D)** Hierarchical organization of nociceptive pathways as revealed by activity mapping.

## Discussion

Understanding the molecular organization of nociceptive responses has been hindered by the lack of tools that can simultaneously resolve both transcriptomic and functional properties. Immediate-early gene-driven labeling has revolutionized CNS ensemble identification^49-51^ but fails in PNS neurons; whereas CaMPARI enables unbiased activity-based ensemble identification in DRG.

Activity labeling revealed a robust domain-specific organization: cutaneous C-HTMRs (*Nefh*^*-*^) were dominated by NP1 (*Mrgprd*) and NP2 (*Mrgpra3/b4*) populations, in line with previous findings in the trigeminal system^34,35^, while visceral C-nociceptors were primarily PEP1 (Adra2a) and, to a lesser degree, PEP1 (Oprk1) neurons^3,52^. Among A-HTMRs (*Nefh*^*+*^), PEP3 subsets segregated into cutaneous (*S100a16*) and visceral (*Adm*) specializations, with a PEP3 subgroup (*Uts2b*^+^) tuned to bladder stretch. This organization was supported by calcium imaging and peripheral tracing, matching emerging anatomical evidence for restricted projection fields^3,13,53^.

Our data demonstrate that noxious stimuli recruit a rich repertoire of nociceptor types that follow a hierarchical transcriptomic organization. The major functional divide reflects critical neurophysiological (myelinated *Nefh*^*+*^ versus unmyelinated *Nefh*^*-*^ conduction) and neurochemical properties (neuropeptide content). However, in line with increasing specificity, broad nociceptor classes (*Bmpr1b*^*+*^ and *Trpv1*^*+*^*) are* subdivided into distinct types associated with specific body domains (through *Adm*^*+*^ and *Adra2a*^*+*^ subtypes) and organs like bladder (through *Uts2b*^*+*^*Adm*^*+*^ cells, Fig. 7D). Taken together, our findings provide a mechanistic basis for the proposed division between interoceptive and exteroceptive nociceptive pathways^4^, demonstrating that this distinction reflects a genetic hierarchy spanning across nociceptor classes. This layered functional-anatomical organization of high-threshold mechanoreceptors (HTMRs) signaling likely reflects a developmental trajectory in which intrinsic genetic programs establish broad neuronal identity, subsequently refined by peripheral target-derived cues^29,30^.

Most research on pain focuses on cutaneous responses, while many clinical pain syndromes involve deep, visceral sensations^4^. Critically, transcriptomic coverage of human sensory neurons is still limited to ganglia innervating the limbs and body trunk, and therefore highly enriched in cutaneous innervation^54,55^. However, our framework may help contextualize future, more extensive datasets from human sacral ganglia within a physiological setting. Although our study focuses on acute noxious stimuli, certain mechano-nociceptors are critical in pathological states like colitis^3^. Conversely, even populations not engaged in the ensemble in baseline conditions may become functionally relevant under pathological conditions in specific organs^34,56^, or mediate reflexive responses^57^. Determining how such pathological states alter ensemble response profiles clearly warrants further investigation.

1. Our system-level analysis provides a blueprint for nociceptive system organization and establishes a generalizable framework for mapping peripheral pathways across the body. The identification of a transcriptomic hierarchy reveals that visceral and cutaneous mechano-nociceptive pathways are fundamentally distinct, outlining the need for increased focus on viscera-specific populations in the development of precision treatments. In summary, our work can facilitate future efforts to dissect and manipulate the peripheral nervous system to understand how naturalistic stimuli shape internal states.

## ACKNOWLEDGMENTS

We are grateful to Prof. Patrik Ernfors for critical reading of the manuscript, and to members of the Ernfors group: Dr. Dmitri Usoskin, Dr. Jussi Kupari, and Dr. Jie Su for thoughtful comments and advice. We thank Prof. Marie Larsson and Dr. Francis Hopkins for advice on scRNA-seq, and Prof. Markus Heilig, Dr. Nima Ghitani, and Dr. Joshua Emrick for helpful discussions. We also thank Bengt Ragnemalm for assistance in device development. We are indebted to the Linköping University Imaging Facility, led by Dr. Vesa Loitto, for access and assistance with anatomical imaging, and to the National Genomics Infrastructure and SciLifeLab Sweden particularly Helena Parra Acero, Anja Mezger, and Michelle Ljungberg for their help and flexibility with scRNA-seq. We also thank the Center for Biomedical Resources at Linköping University, especially Emma Egedal, Monika Kozak-Ljunggren, and Emelie Schelin, for professional maintenance of the animal colony and Dr. Joost Wiskerke at Neuroimaging Platform for assistance with animal surgeries. We are grateful to Batuhan Yagan and Simon Ekborn for help with tissue collection. We thank Prof. David Ginty for providing the Bmpr1b^Cre^ mouse line. This work was supported by the Swedish Research Council (Vetenskapsrådet 2024-02781), Knut and Alice Wallenberg Foundation Fellowship, and the Brain Foundation (Hjarnfonden 2025-00169) grants to M.S., as well as by a Wallenberg Project Grant 2019.0047 to H.O. and M.L.

## AUTHOR CONTRIBUTIONS

Conceptualization - FdF, ML, MS, Data Curation - FdF, GBC, MBMC, IS, MB, MS, Formal analysis - FdF, GBC, MBMC, IS, MB, MS, Funding Acquisiton - HO, ML, MS, Investigation - FdF, GBC, MB, CK, LFN, ML, Software - MBMC, IS, MS, Supervision – MS, Validation - FdF, GBC, MBMC, MS, Visualization - FdF, GBC, MBMC, MS, Writing - original draft – MS, Writing - review & editing - FdF, GBC, MBMC, IS, MB, CK, LFN, HO, ML, MS

