## Supplementary information for "Hierarchical organization of mechano-nociceptive pathways revealed by activity labeling"

**Supplementary materials tables and figures.**
**Data Tables:**
**Supplementary table 1.** Cell counts mapped to general categories for tail brush, perianal brush, tail pinch and colorectal stroke related to Fig. 3.
**Supplementary table 2.** Pairwise comparisons statistics for post hoc analysis of defined subgroups within tested stimuli related to Fig. 3D
**Supplementary table 3** Pairwise comparisons statistics for post hoc analysis of defined subgroups within tested stimuli within myelination classes related to Fig. 3E.
**Supplementary table 4** Cell counts mapped to specified transcriptomic categories for Tail pinch, hair-pull, anorectal distension stimuli in pan-neuronal CaMPARI mice and colorectal distension and bladder stretch in $Trpv1^{lin}$:CaMPARI, related to Fig. 4 and Fig. 6.
**Supplementary table 5.** Pairwise comparison of response pattern of C-nociceptors to high intensity stimulation related to Fig. 4C.
**Supplementary table 6** Post hoc comparisons of the extent of transcriptomic classes activated by different stimuli. Related to Figure 4D.
**Supplementary table 7** Pairwise comparison of response pattern of A-nociceptors to high intensity stimulation related to Figure 6C.
**Supplementary table 8** Post hoc pairwise comparisons of the extent of transcriptomic classes activated by different stimuli. Related to Figure 6D.
Figures:
**Supplementary Figure 1.1** Conversion of active cells in the L6 DRG by brushing
**Supplementary Figure 1.2** Example conversions of a range of gentle and noxious cutaneous stimuli
**Supplementary Figure 2.1** Activation of sensory neurons by cutaneous and visceral stimuli
**Supplementary Figure 2.2** Detection of functional modules within pelvic sensory neurons
**Supplementary Figure 2.3** Single-cell RNA sequencing cellular atlas preprocessing and quality control
**Supplementary Figure 2.4** Cluster characteristic gene expression matrix
**Supplementary Figure 2.5** Expression analysis of selected marker genes
**Supplementary Figure 2.6** Comparison of pelvic sensory neuron atlas with published DRG datasets
**Supplementary Figure 2.7** Distribution of *Bmpr1b*, *Adm,* and *Adra2a* neurons along the neuroaxis
**Supplementary Figure 3.1** Ensemble representation of the Tail brush stimulus
**Supplementary Figure 4.1** Example conversions of noxious visceral stimuli
**Supplementary Figure 4.2** C-nociceptors ensemble composition for Hair pulling and Anorectal distension
**Supplementary Figure 4.3.** Peripheral viral labeling of pelvic DRG neurons
**Supplementary Figure 5.1** Activation of myelinated sensory neurons by cutaneous and visceral stimuli
**Supplementary Figure 5.2** Detection of functional cellular modules within myelinated neurons
**Supplementary Figure 6.1** A-nociceptors ensemble composition for Hair pulling and Anorectal distension

**Supplementary table 1.** Cell counts mapped to specified category for tail brush (N=2), perianal brush (N=6), tail pinch (N=12) and colorectal stroke stimuli (N=15 mice).

| | A LTMRs | A nociceptors | C LTMRs | C nociceptors |
|---|---|---|---|---|
| Tail Brush | 16 | 1 | 29 | 6 |
| Perianal brush | 39 | 2 | 32 | 16 |
| Tail Pinch | 33 | 20 | 41 | 43 |
| Colorectal stroke | 41 | 31 | 10 | 42 |

**Supplementary table 2.** Pairwise comparisons statistics for post hoc analysis of defined subgroups within tested stimuli related to Fig. 3D.

| Stimuli 1 | Stimuli 2 | Group | G-test statistics | Multiple comparison corrected p-value (Holm method) |
|---|---|---|---|---|
| Tail Brush | Tail Pinch | A LTMRs | 3.58E-01 | 1.00E+00 |
| Tail Brush | Tail Pinch | A nociceptors | 9.92E-03 | 9.92E-02 |
| Tail Brush | Tail Pinch | C LTMRs | 1.34E-03 | 2.02E-02 |
| Tail Brush | Tail Pinch | C nociceptors | 5.16E-03 | 5.67E-02 |
| Tail Brush | Stroke | A LTMRs | 8.60E-01 | 8.60E-01 |
| Tail Brush | Stroke | A nociceptors | 8.17E-05 | 1.31E-03 |
| Tail Brush | Stroke | C LTMRs | 3.18E-11 | 6.37E-10 |
| Tail Brush | Stroke | C nociceptors | 2.69E-03 | 3.23E-02 |
| Perianal brush | Tail Pinch | A LTMRs | 2.21E-03 | 2.87E-02 |
| Perianal brush | Tail Pinch | A nociceptors | 2.14E-03 | 3.00E-02 |
| Perianal brush | Tail Pinch | C LTMRs | 3.83E-01 | 1.00E+00 |
| Perianal brush | Tail Pinch | C nociceptors | 2.99E-02 | 2.39E-01 |
| Perianal brush | Stroke | A LTMRs | 1.17E-01 | 7.02E-01 |
| Perianal brush | Stroke | A nociceptors | 1.56E-06 | 2.81E-05 |
| Perianal brush | Stroke | C LTMRs | 5.67E-07 | 1.08E-05 |
| Perianal brush | Stroke | C nociceptors | 1.23E-02 | 1.11E-01 |
| Tail Pinch | Stroke | A LTMRs | 1.30E-01 | 6.52E-01 |
| Tail Pinch | Stroke | A nociceptors | 4.20E-02 | 2.94E-01 |
| Tail Pinch | Stroke | C LTMRs | 7.99E-06 | 1.36E-04 |
| Tail Pinch | Stroke | C nociceptors | 6.93E-01 | 1.00E+00 |

**Supplementary table 3** Pairwise comparisons statistics for post hoc analysis of defined subgroups within tested stimuli within myelination classes related to Fig. 3E.

| Stimulus 1 | Stimulus 2 | Group | Fisher exact test. | Multiple comparison corrected p-value (Holm method) |
|---|---|---|---|---|
| Tail Brush | Tail Pinch | C nociceptors | 8.76E-04 | 8.76E-04 |
| Tail Brush | CR stroke | C nociceptors | 7.59E-09 | 6.07E-08 |
| Per. brush | CR stroke | C nociceptors | 2.05E-06 | 1.23E-05 |
| Tail Pinch | CR stroke | C nociceptors | 5.37E-04 | 1.61E-03 |
| Tail Brush | Tail Pinch | A nociceptors | 1.43E-02 | 2.86E-02 |
| Tail Brush | CR stroke | A nociceptors | 4.08E-03 | 1.22E-02 |
| Perianal brush | Tail Pinch | A nociceptors | 1.55E-04 | 9.29E-04 |
| Perianal brush | CR stroke | A nociceptors | 8.83E-06 | 7.07E-05 |

**Supplementary table 4** Cell counts mapped to specified transcriptomic categories for Tail pinch (N=12), hair-pull (N=4), anorectal distension (N=9), colorectal stroke (N=15 mice) stimuli in pan-neuronal CaMPARI mice and colorectal distension (N=8) and bladder stretch (N=6) in $Trpv1^{lin};^{R26LSL-CaMPARI}$. Related to Fig.4 and Fig. 6.

| | Tail Pinch | Hair Pull | Anorectal distension | Colorectal stroke | Colorectal distension | Bladder distension |
|---|---|---|---|---|---|---|
| NP1 (Mrgprd) | 23 | 13 | 24 | 8 | 12 | 8 |
| NP2 (MRGPRA3/B4) | 7 | 2 | 6 | 0 | 2 | 1 |
| NP3 (Sst) | 3 | 1 | 2 | 2 | 2 | 0 |
| PEP1 (Sstr2) | 2 | 1 | 7 | 6 | 0 | 2 |
| PEP1 (Adra2a) | 1 | 0 | 2 | 12 | 11 | 16 |
| PEP1 (Oprk1) | 7 | 2 | 4 | 14 | 11 | 6 |
| C-LTMR (Th) | 41 | 66 | 34 | 10 | 33 | 6 |
| PEP3 (S100a16) | 20 | 5 | 1 | 9 | 2 | 1 |
| PEP3 (Adm) | 0 | 1 | 13 | 22 | 11 | 33 |
| PEP2 (Smr2) | 0 | 1 | 0 | 0 | 0 | 2 |
| Ab-LTMR (Calb1) | 11 | 1 | 2 | 4 | 1 | 3 |
| Ad-LTMR (Ntrk2h) | 22 | 26 | 17 | 24 | 0 | 6 |
| Prop. (Pvalb) | 0 | 4 | 5 | 13 | 0 | 3 |
| Unassigned | 5 | 4 | 1 | 1 | 3 | 1 |

**Supplementary table 5.** Pairwise comparison of response pattern of C-nociceptors to high intensity stimulation related to Fig. 4C.

| Stimulus 1 | Stimulus 2 | G-test statistics | Calculated p-value | Multiple comparison corrected p-value (Holm method) |
|---|---|---|---|---|
| Tail Pinch | Hair pull | 1.890 | 8.64E-01 | 8.64E-01 |
| Tail Pinch | Anorectal distension | 4.213 | 5.19E-01 | 1.00E+00 |
| Tail Pinch | CR stroke | 30.743 | 1.05E-05 | 6.32E-05 |
| Hair pull | Anorectal distension | 3.393 | 6.40E-01 | 1.00E+00 |
| Hair pull | CR stroke | 26.146 | 8.36E-05 | 3.34E-04 |
| Anorectal distension | CR stroke | 30.472 | 1.19E-05 | 5.95E-05 |

**Supplementary table 6.** Post hoc pairwise comparisons of the extent of transcriptomic classes activated by different stimuli. Related to Figure 4D.

| Stimulus 1 | Stimulus 2 | Cell class | Fisher exact test. | Multiple comparison corrected p-value (Holm method) |
|---|---|---|---|---|
| Hair pull | CR stroke | NP1 (Mrgprd) | 3.35E-04 | 6.02E-03 |
| Tail Pinch | CR stroke | NP1 (Mrgprd) | 1.35E-03 | 2.30E-02 |
| Anorectal distension | CR stroke | NP1 (Mrgprd) | 1.63E-03 | 2.45E-02 |
| Tail Pinch | CR stroke | PEP1 (Adra2a) | 1.57E-03 | 2.51E-02 |
| Anorectal distension | CR stroke | PEP1 (Adra2a) | 2.84E-03 | 3.98E-02 |
| Anorectal distension | CR stroke | PEP1 (Oprk1) | 7.29E-03 | 9.47E-02 |
| Tail Pinch | CR stroke | NP2 (Mrgpra3/b4) | 1.19E-02 | 1.31E-01 |
| Hair pull | CR stroke | PEP1 (Adra2a) | 1.19E-02 | 1.42E-01 |
| Anorectal distension | CR stroke | NP2 (Mrgpra3/b4) | 2.65E-02 | 2.65E-01 |
| Hair pull | CR stroke | NP2 (Mrgpra3/b4) | 9.34E-02 | 8.41E-01 |
| Tail Pinch | CR stroke | PEP1 (Oprk1) | 1.29E-01 | 9.06E-01 |
| Hair pull | CR stroke | PEP1 (Oprk1) | 1.14E-01 | 9.09E-01 |
| Tail Pinch | CR stroke | NP3 (Sst) | 6.76E-01 | 1.00E+00 |
| Tail Pinch | CR stroke | PEP1 (Sstr2) | 2.65E-01 | 1.00E+00 |
| Hair pull | CR stroke | NP3 (Sst) | 1.00E+00 | 1.00E+00 |
| Hair pull | CR stroke | PEP1 (Sstr2) | 4.18E-01 | 1.00E+00 |
| Anorectal distension | CR stroke | NP3 (Sst) | 1.00E+00 | 1.00E+00 |
| Anorectal distension | CR stroke | PEP1 (Sstr2) | 1.00E+00 | 1.00E+00 |

**Supplementary table 7.** Pairwise comparison of response pattern of A-nociceptors to high intensity stimulation related to Figure 6C.

| Stimulus 1 | Stimulus 2 | G-test statistics | Calculated p-value | Multiple comparison corrected p-value (Holm method) |
|---|---|---|---|---|
| Tail Pinch | Hair pull | 5.883 | 5.28E-02 | 1.06E-01 |
| Tail Pinch | CR stroke | 32.386 | 9.28E-08 | 4.64E-07 |
| Tail Pinch | Anorectal distension | 38.029 | 5.52E-09 | 3.31E-08 |
| Hair pull | CR stroke | 9.831 | 7.33E-03 | 2.20E-02 |
| Hair pull | Anorectal distension | 14.122 | 8.58E-04 | 3.43E-03 |
| CR stroke | Anorectal distension | 3.117 | 2.10E-01 | 2.10E-01 |

**Supplementary table 8.** Post hoc pairwise comparisons of the extent of transcriptomic classes activated by different stimuli. Related to Figure 6D.

| Stimulus 1 | Stimulus 2 | Cell class | Fisher exact test. | Multiple comparison corrected p-value (Holm method) |
|---|---|---|---|---|
| Tail Pinch | CR stroke | PEP3 (Adm) | 1.68E-07 | 1.51E-06 |
| Tail Pinch | CR stroke | PEP3 (S100a16) | 1.68E-07 | 1.68E-06 |
| Tail Pinch | CR stroke | PEP2 (Smr2) | 1.00E+00 | 1.00E+00 |
| Tail Pinch | Anorectal distension | PEP3 (Adm) | 1.51E-08 | 1.73E-07 |
| Tail Pinch | Anorectal distension | PEP3 (S100a16) | 1.51E-08 | 1.73E-07 |
| Tail Pinch | Anorectal distension | PEP2 (Smr2) | 1.00E+00 | 1.00E+00 |
| Hair pull | CR stroke | PEP3 (Adm) | 9.63E-03 | 5.78E-02 |
| Hair pull | CR stroke | PEP3 (S100a16) | 7.72E-02 | 3.86E-01 |
| Hair pull | CR stroke | PEP2 (Smr2) | 1.84E-01 | 7.37E-01 |
| Hair pull | Anorectal distension | PEP3 (Adm) | 8.51E-04 | 6.81E-03 |
| Hair pull | Anorectal distension | PEP3 (S100a16) | 5.55E-03 | 3.88E-02 |
| Hair pull | Anorectal distension | PEP2 (Smr2) | 3.33E-01 | 1.00E+00 |

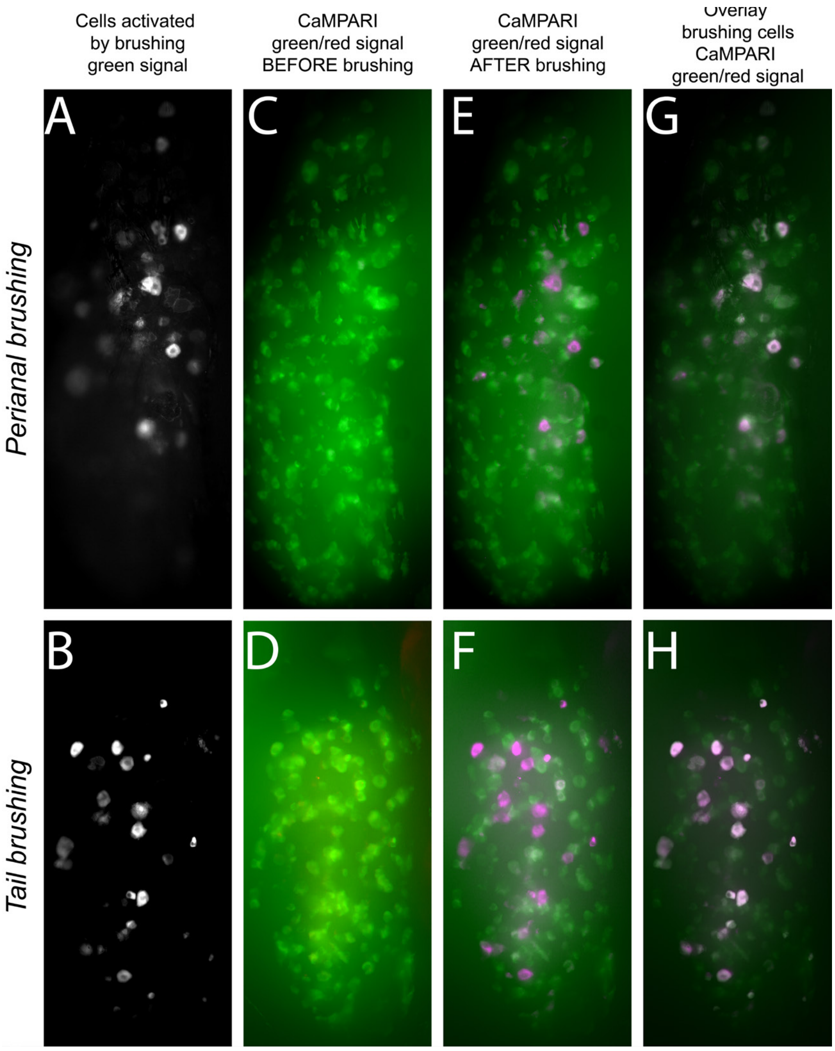


**Supplementary Figure 1.1 Conversion of active cells in the L6 DRG by brushing A-B)** CaMPARI, acting as a reverse calcium indicator, reports activity by dimming of basal green fluorescence in activated cells. Standard deviation projections of green fluorescence during stimulation visualize activated neurons in L6 DRG during perianal brushing (A) and tail brushing (B). **C–F)** Overlay of CaMPARI fluorescence in live L6 DRG, showing the basal form (green, emission λ = 520 nm) and the photoconverted form (red, emission λ = 580 nm, pseudocolored magenta). Panels C-D show pre-stimulus (before brushing), and panels **E-F** show post-stimulus (after brushing). **G-H)** Overlay of cells identified by acute activation during brushing through fluorescence quenching with cells photoconverted by coupling stimulation with UV illumination, showing overlap between functional activity readout and CaMPARI conversion. Look up tables (LUTs) for green and magenta are set independently for each preparation due to differences in e.g. out of focus fluorescence levels).

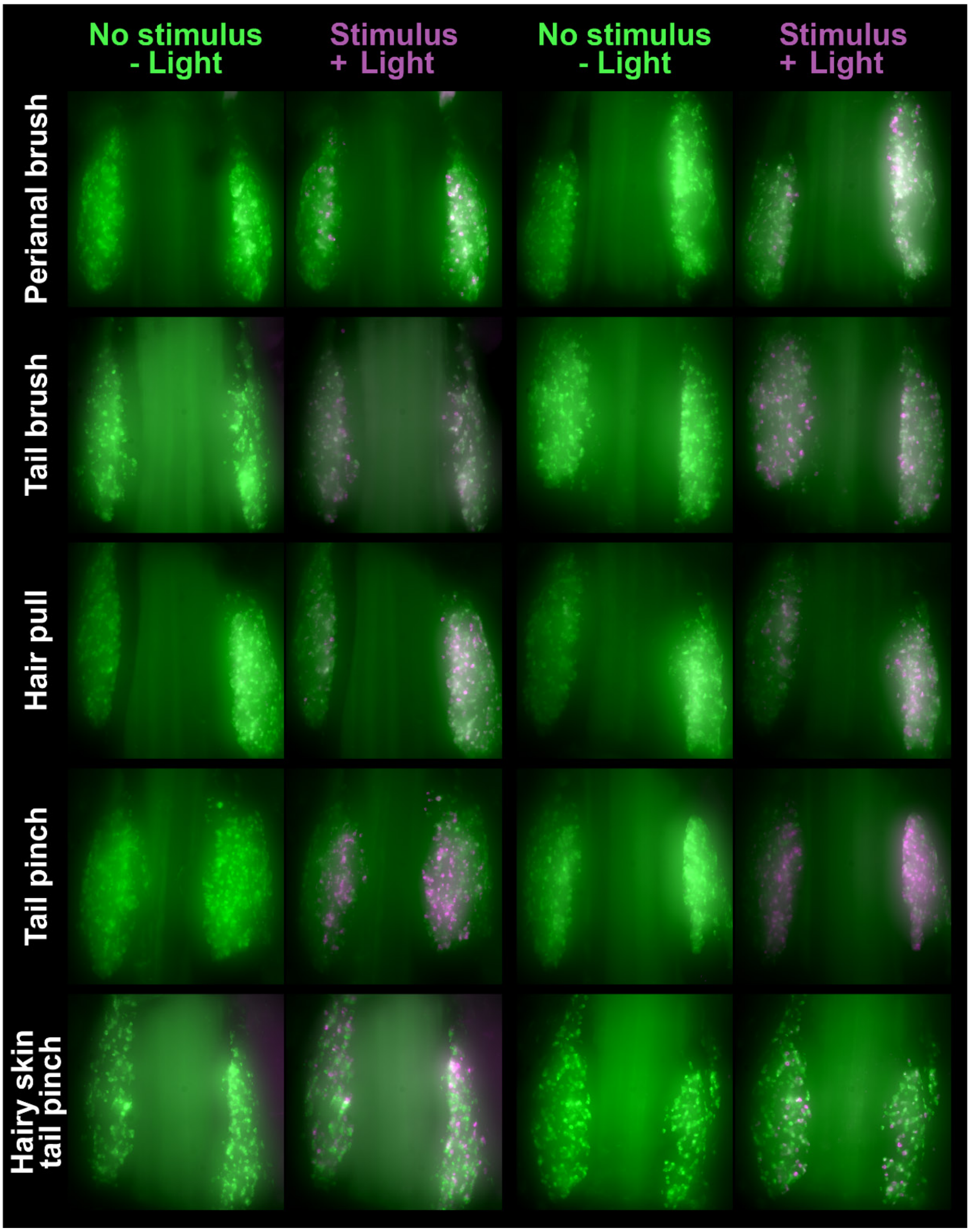


**Supplementary Figure 1.2 Example conversions of a range of gentle and noxious cutaneous stimuli** Overlays of CaMPARI fluorescence in live DRG, showing the basal form (green, emission λ = 520 nm) and the photoconverted form (red, emission λ = 580 nm, pseudocolored magenta) before and after stimuli conversion. Two representative preparations are shown for each stimulus, for a range of gentle and noxious cutaneous stimulations. Look up tables (LUTs) for green and magenta are set independently for each preparation due to differences in e.g out of focus fluorescence levels). Note 'no stimulus' images have been acquired after a mock UV stimulation.

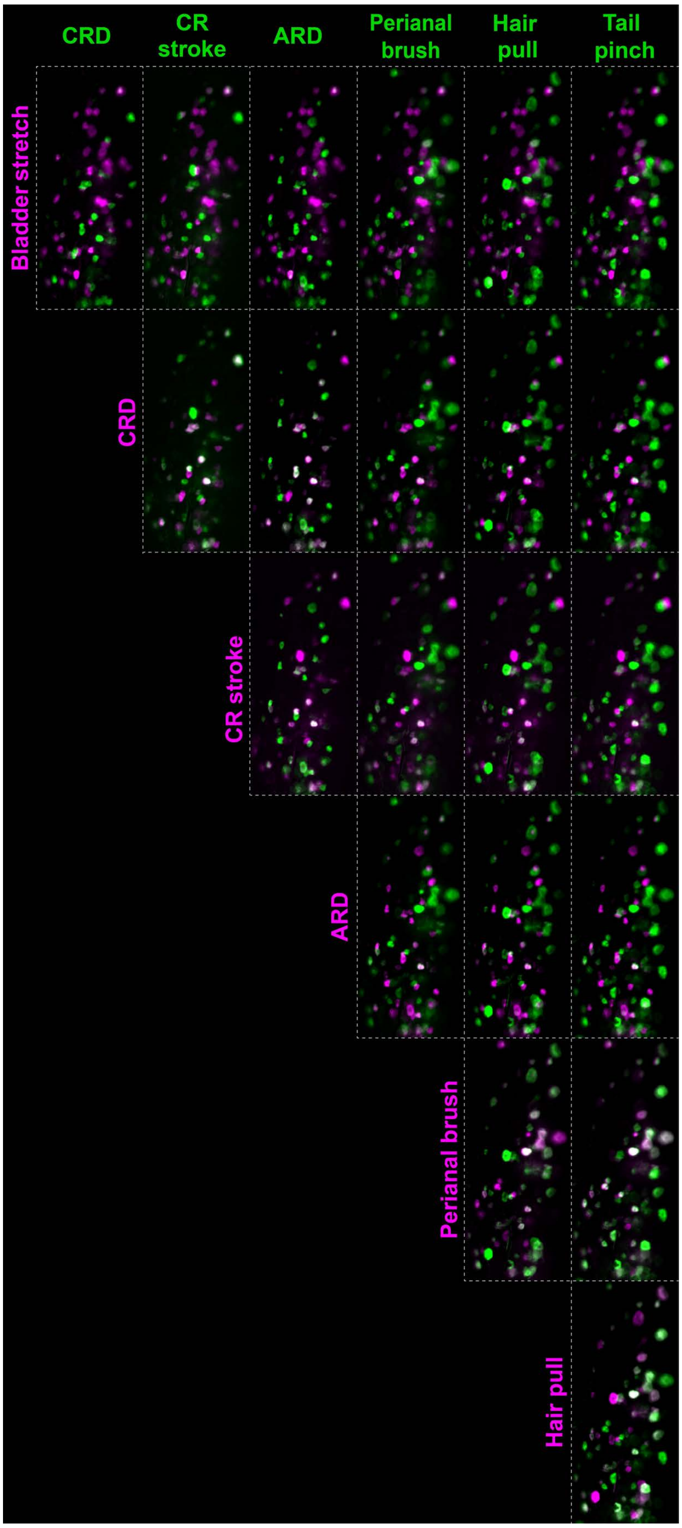


**Supplementary Figure 2.1 Activation of sensory neurons by cutaneous and visceral stimuli** Responses recorded in the mosaic GCaMP6f model. Each panel shows deviation projection visualizing cells activated by the stimulus indicated in the row (magenta) and the stimulus indicated in the column (green). Overlap (white) marks neurons activated by both stimuli. The stimulus panel included bladder stretch, colorectal distension (CRD, 80 mmHg), colorectal stroke (CR stroke), anorectal distension (ARD, 80 mmHg), perianal brushing, hair pull, and tail pinch.

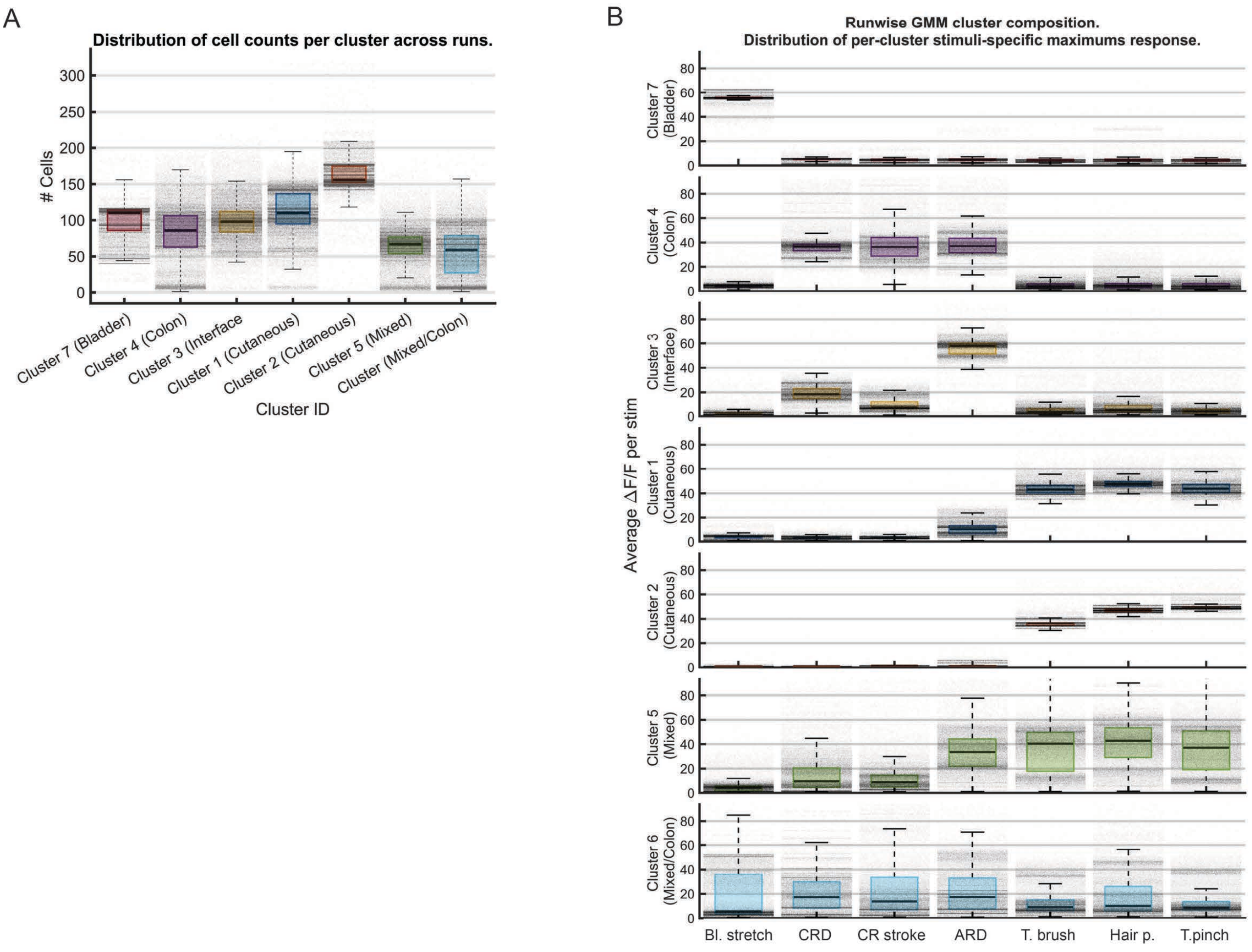


**Supplementary Figure 2.2 Detection of functional modules within pelvic sensory neurons A)** Comparison of Gaussian mixed model (GMM) derived clusters estimated over 50 000 algorithm realizations, graph shows average number of cells qualified into corresponding clusters. **B)** Stability of GMM cluster composition across 50 000 realizations. Box plots show the average stimulus response of cells within each cluster across all realizations; individual dots represent the mean response from a single clustering run. The most representative run was selected for further analysis in Figure 2.

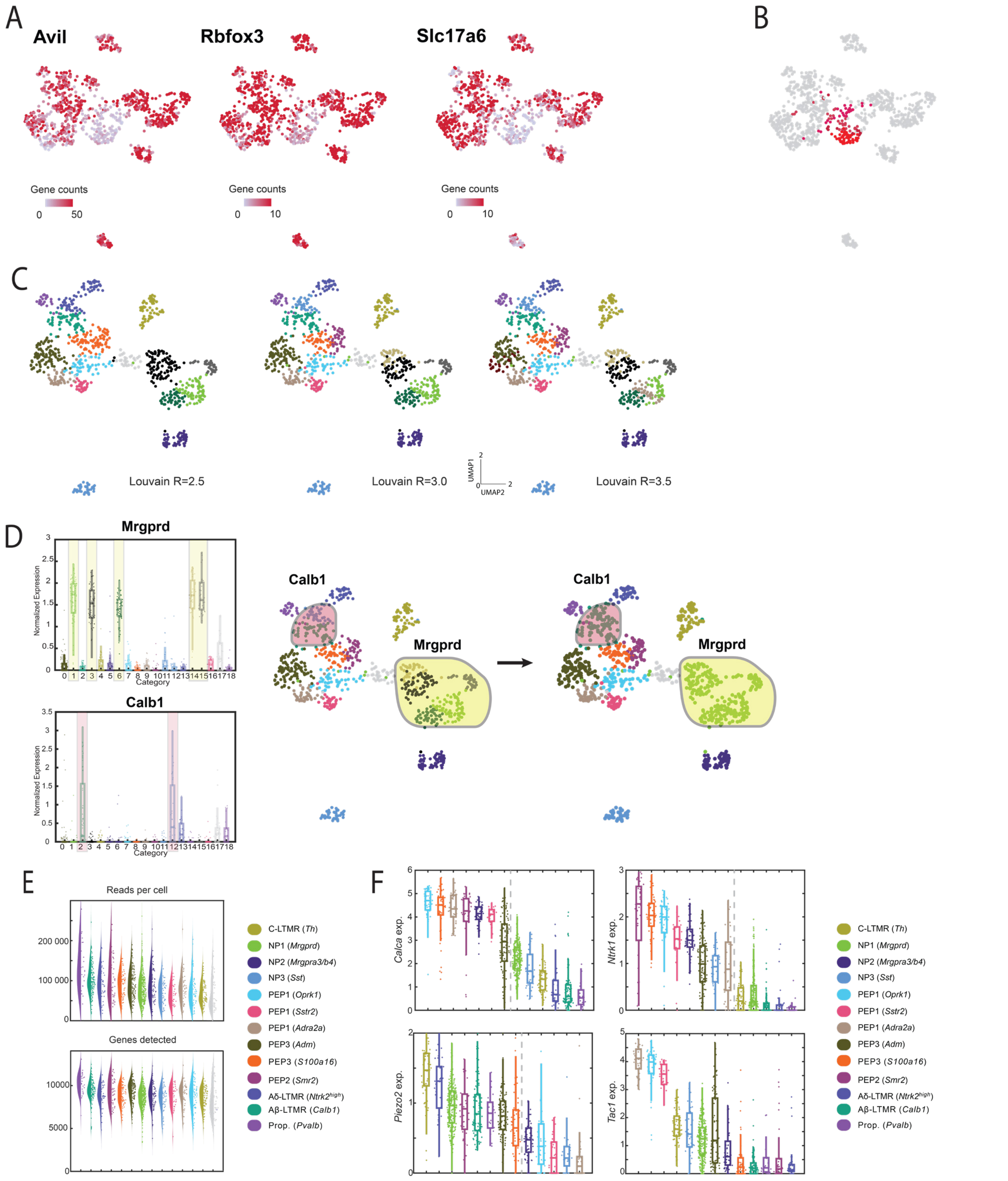


**Supplementary Figure 2.3. Single-cell RNA sequencing cellular atlas preprocessing and quality control** Legend continued on next page…

**… A–B)** Detection of clusters containing low-quality cells. After excluding cells with high mitochondrial gene content (>5%), cells were initially clustered and expression of common markers of primary sensory neurons (*Avil, Rbfox3, Slc17a6*) was assessed. Clusters with low expression of these marker genes were excluded from subsequent analysis. **C)** Clusters detected in the final atlas show little dependence on the clustering resolution parameter (R). **D)** Identification of related clusters for merging. The NP1 (*Mrgprd*) cluster was identified as fragmented based on shared *Mrgprd* expression, and the Aβ-LTMR (*Calb1*) cluster was identified based on shared *Calb1* expression. **E)** Characterization of sequencing depth: distribution of UMI counts per cell within each cluster (top) and number of detected genes (bottom). **F)** Differential expression analysis of representative characteristic and functionally relevant genes across detected clusters.

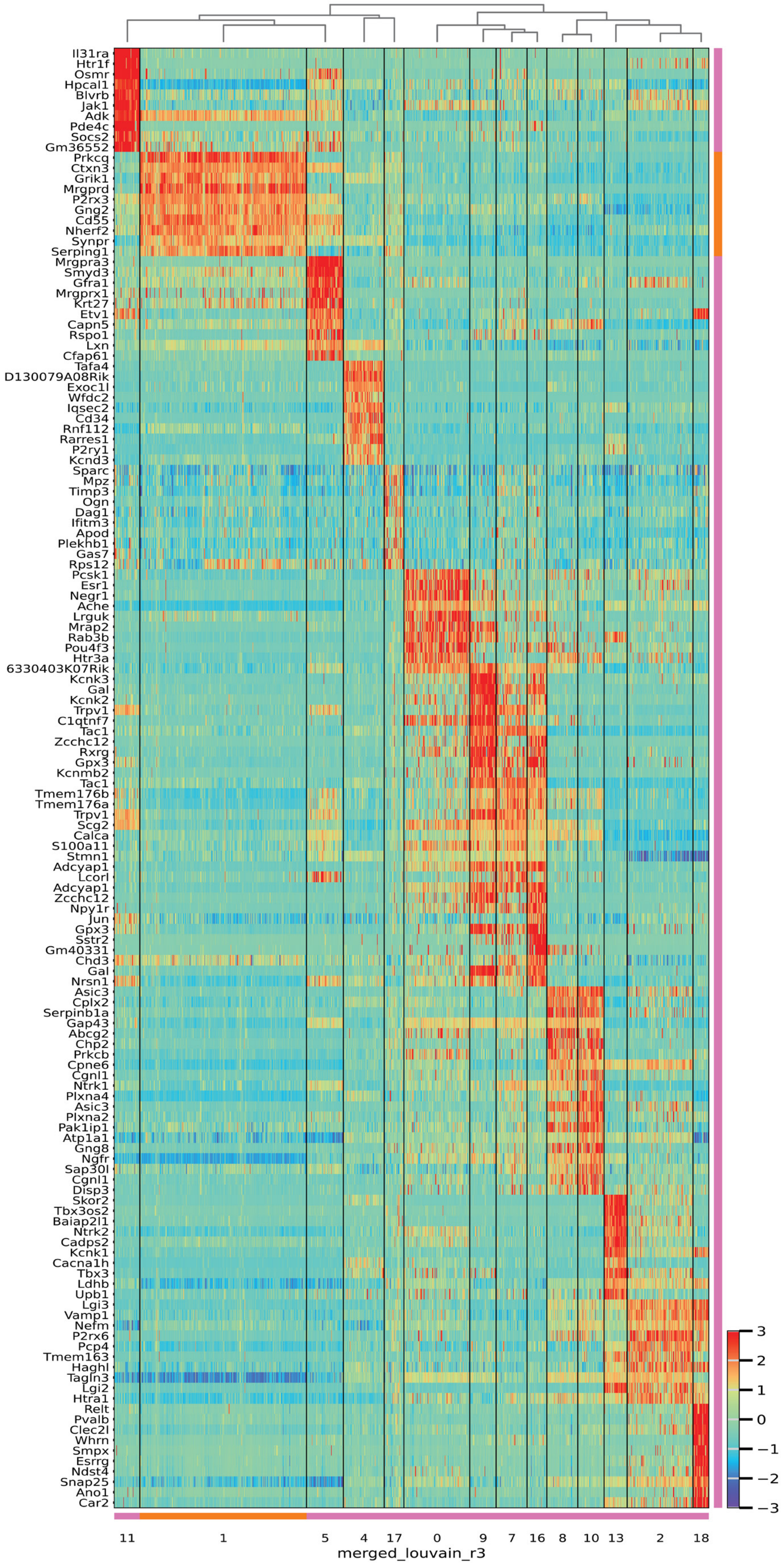


**Supplementary Figure 2.4. Cluster characteristic gene expression matrix** Heatmap showing the expression of ten representative characteristic genes for each cluster.

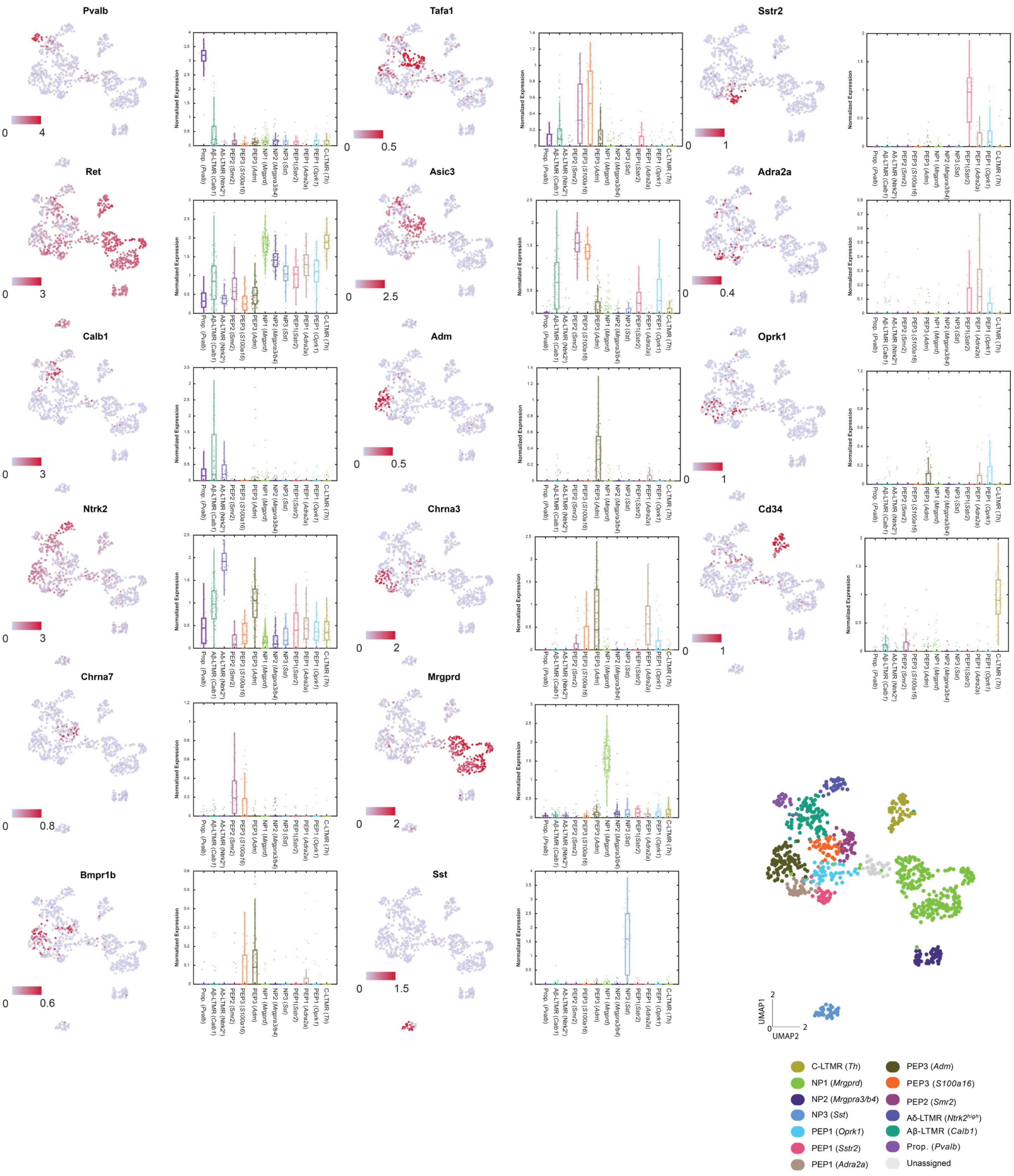


**Supplementary Figure 2.5 Expression analysis of selected marker genes** UMAP and bar graph representation of characteristic markers used in clusters identification. Scale indicates normalized log-transformed expression levels.

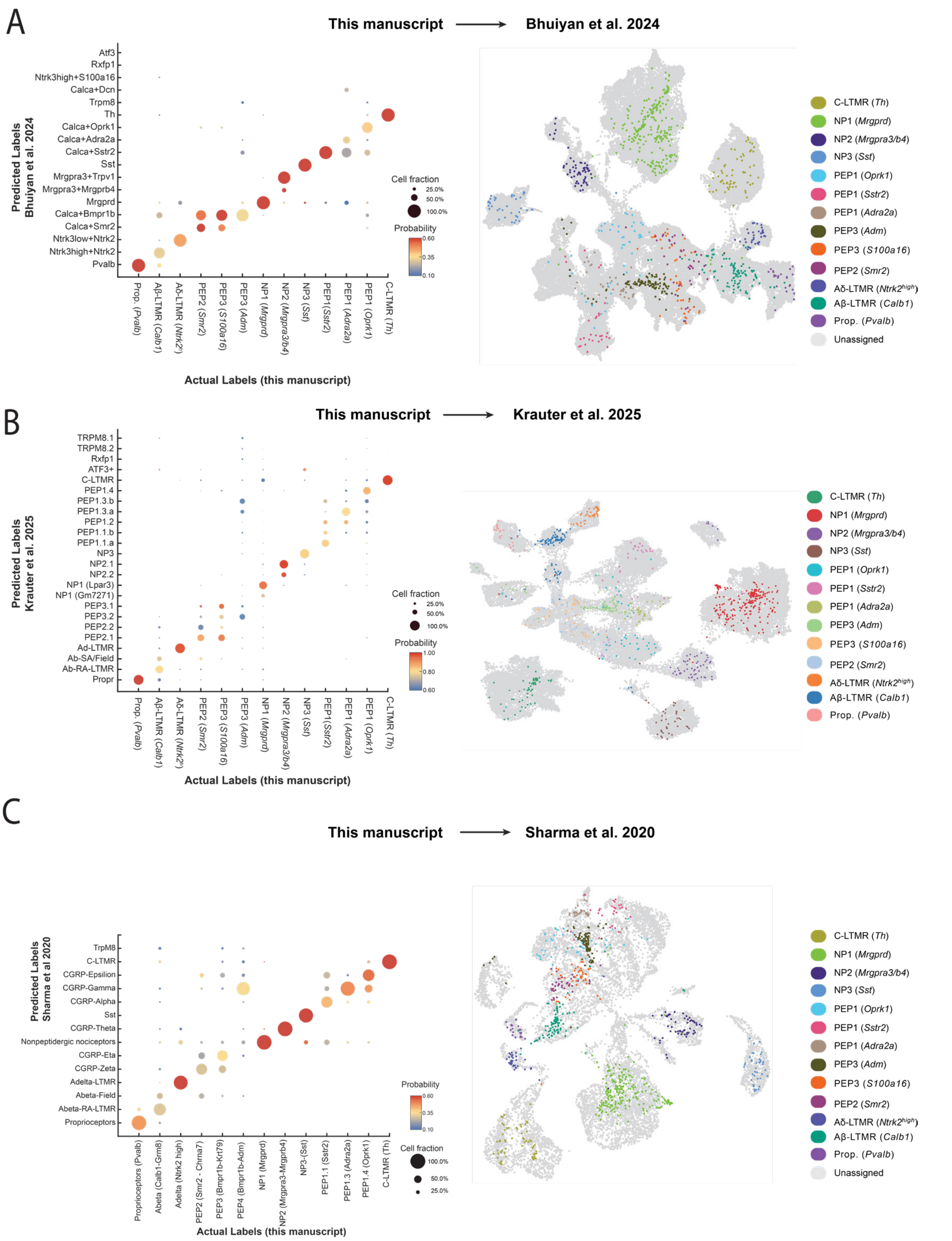


**Supplementary Figure 2.6 Comparison of pelvic sensory neuron atlas with published DRG datasets** Embedding of scRNA-seq pelvic neuron data into general DRG atlases from (A) Bhuiyan et al., 2024; (B) Krauter et al., 2025; and (C) Sharma et al., 2020. Left: bubble plots showing transferred cluster assignments, with labels from the reference atlases mapped onto the pelvic neuron dataset to infer class correspondence. Right: embeddings of pelvic DRG cells into the reference atlases from the source datasets.

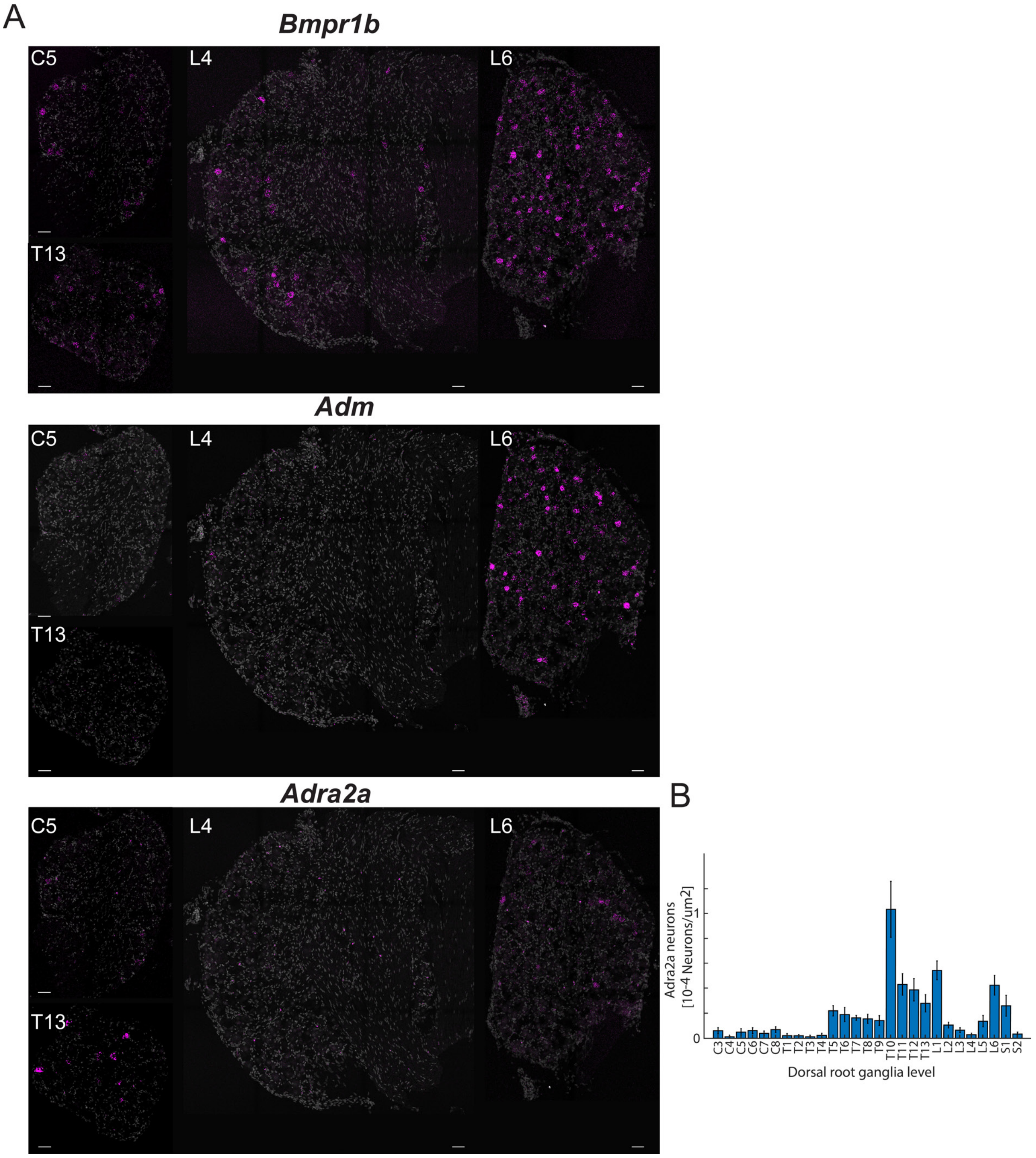


**Supplementary Figure 2.7 Distribution of Bmpr1b, Adm, and Adra2a neurons along the neuroaxis** Related to Figure 2K–L. A) Example FISH images from cervical, thoracic, and lumbar DRG showing expression of Bmpr1b, Adm, and Adra2a. B) Quantification of Adra2a$^+$ neuron abundance along the neuroaxis. Consistent with previous reports (Qi et al., 2024), Adra2a$^+$ neurons were enriched in lower thoracic as well as upper and lower lumbar DRG, but were largely absent from mid-lumbar ganglia innervating the extremities (n ≥ 6; scale bars, 50μm).

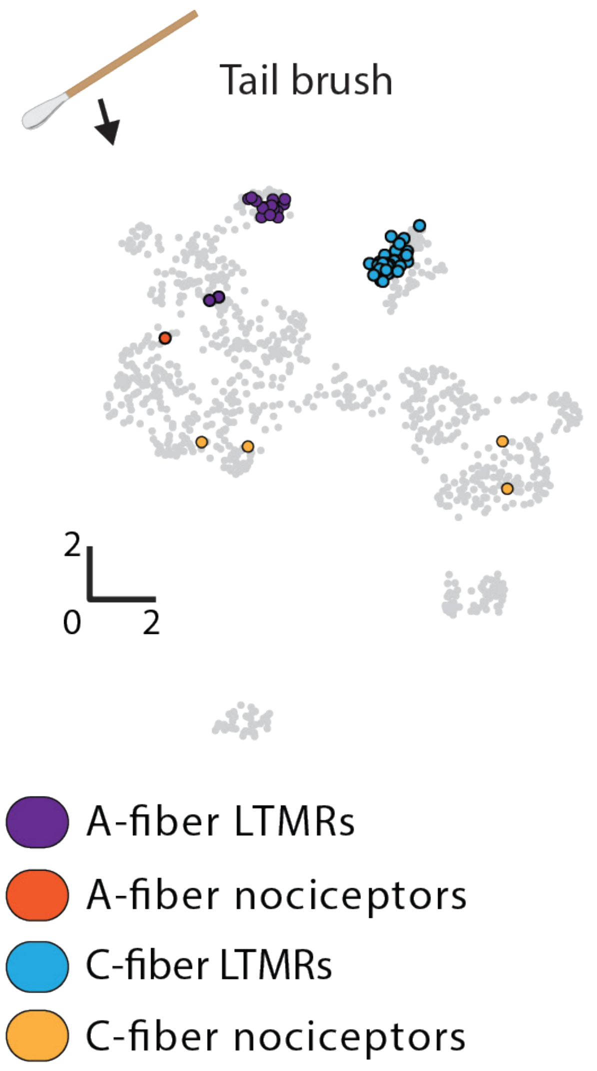


**Supplementary Figure 3.1 Ensemble representation of the Tail brush stimulus** Overlay of the pelvic DRG transcriptomic atlas (gray) with labeled tail brush neurons, color-coded by their class membership.

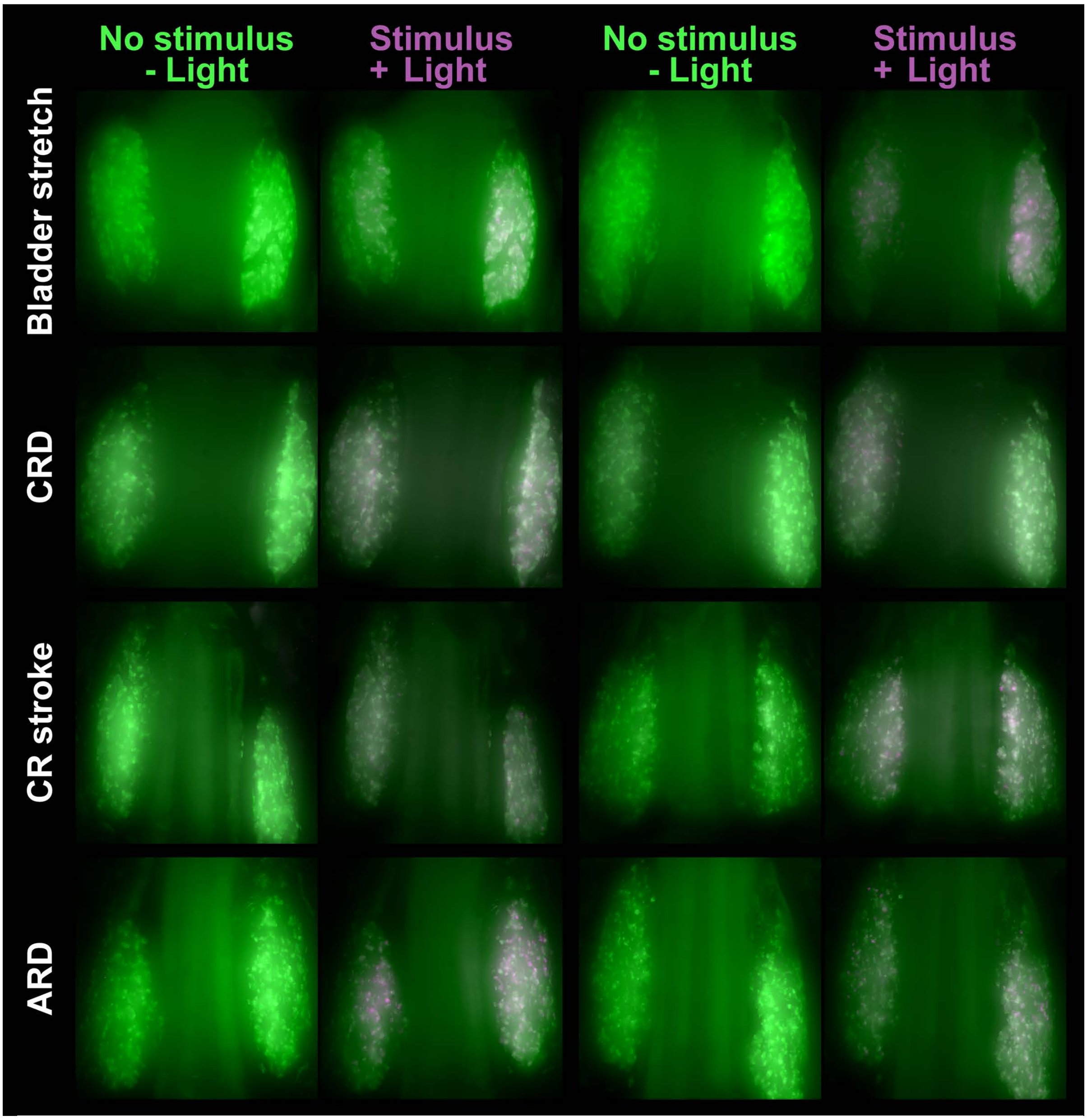


**Supplementary Figure 4.1 Example conversions of noxious visceral stimuli** Overlays of CaMPARI fluorescence in live DRG, showing the basal form (green, emission $\lambda$ = 520 nm) and the photoconverted form (red, emission $\lambda$ = 580 nm, pseudocolored magenta) before and after stimuli conversion. Two representative preparations are shown for each stimulus, for a range visceral stimulus. Look up tables (LUTs) for green and magenta are set independently for each preparation due to differences in e.g. out of focus fluorescence levels).

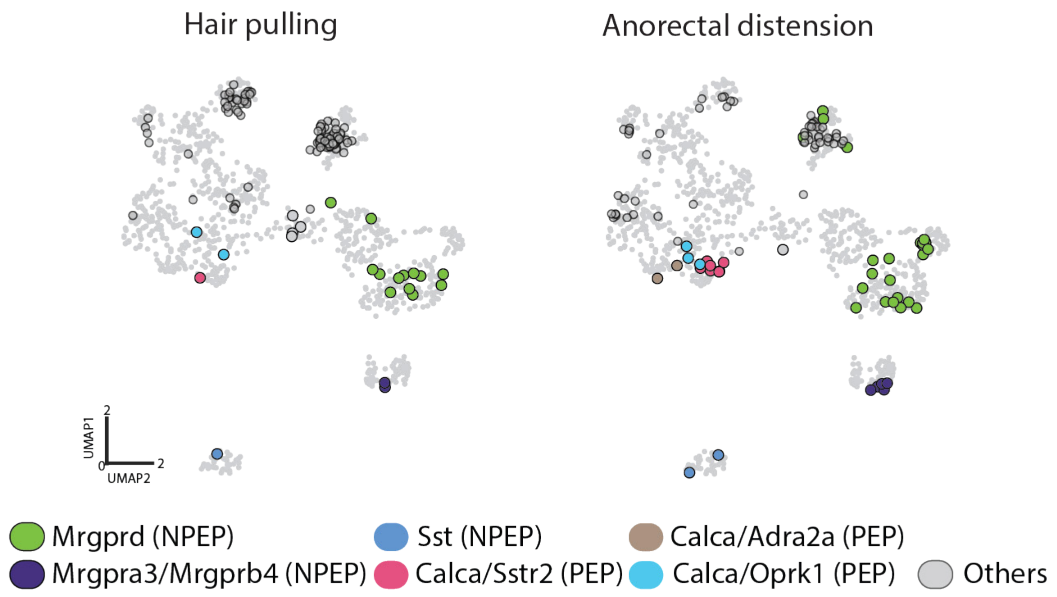


**Supplementary Figure 4.2 C-nociceptors ensemble composition for hair pulling and anorectal distension** Overlay of the pelvic DRG transcriptomic atlas (gray) with stimulus-responsive C-nociceptors, color-coded by their class membership.

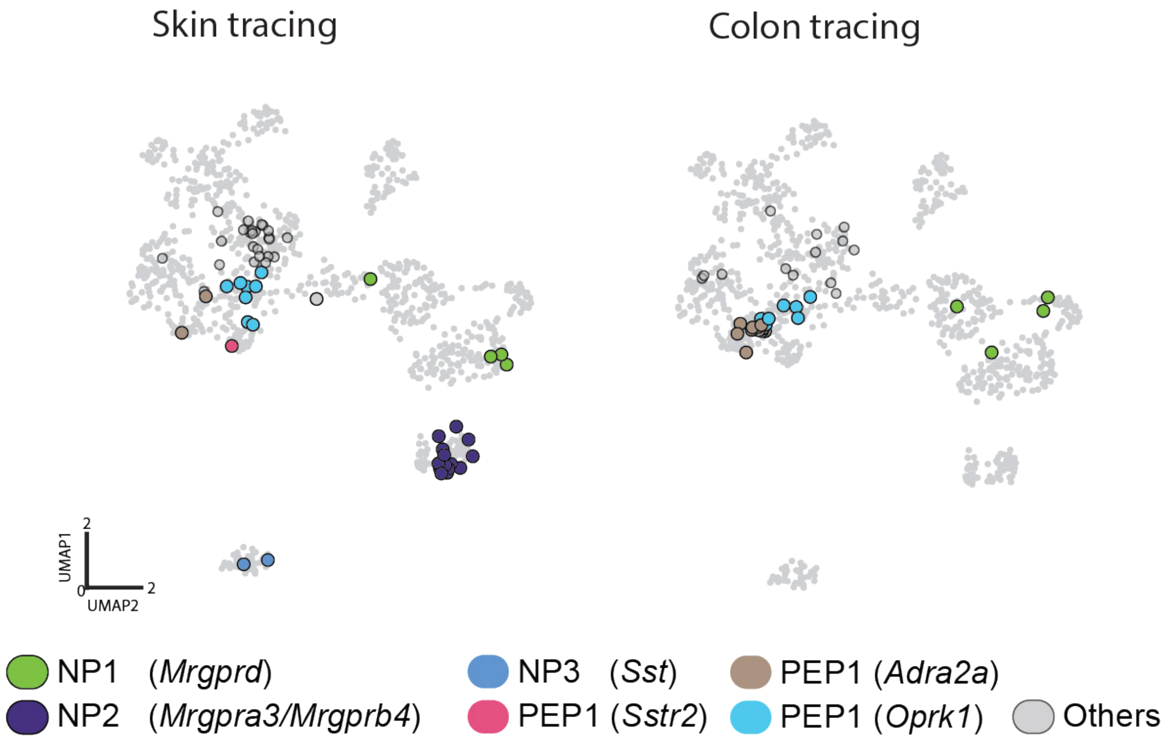


**Supplementary Figure 4.3. Peripheral viral labeling of pelvic DRG neurons** Overlay of the pelvic DRG transcriptomic atlas (gray) with cells labeled by peripheral viral injection, color-coded according to their transcriptomic class membership.

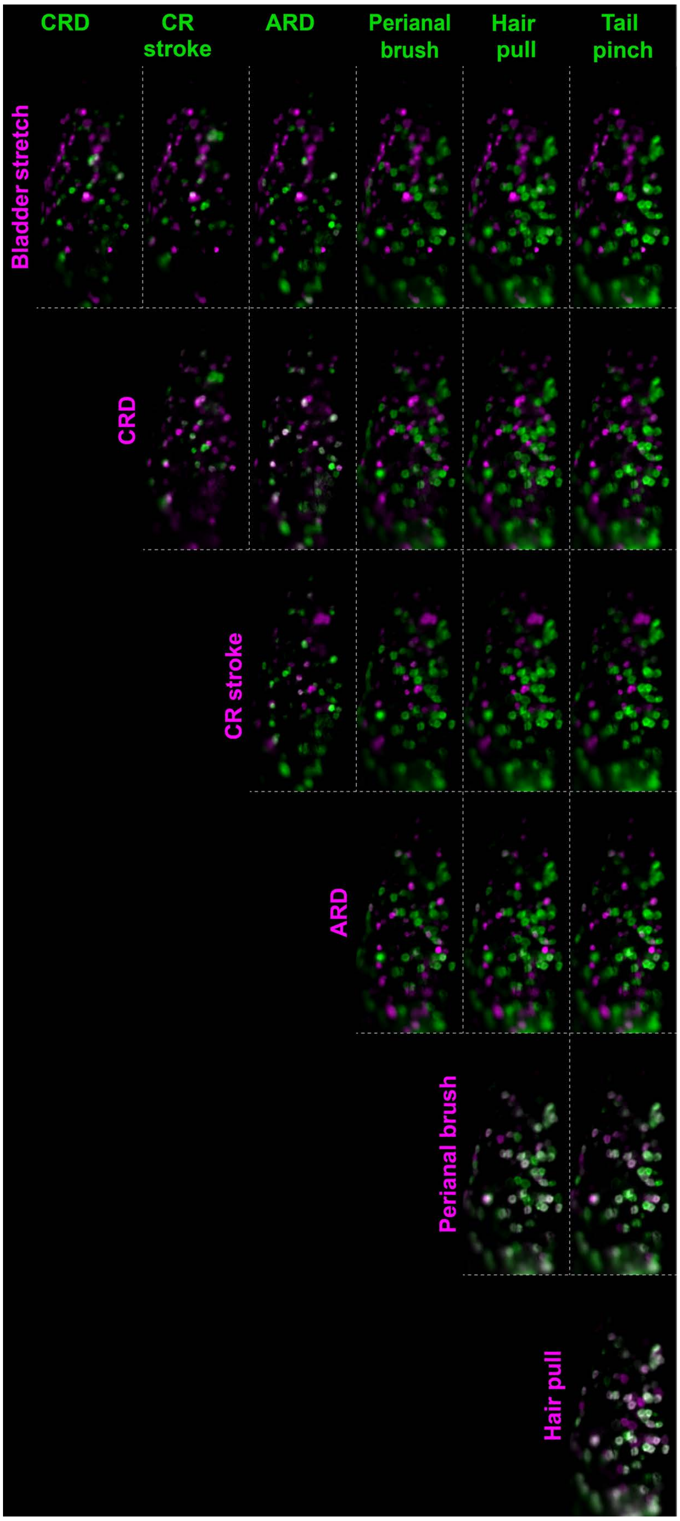


**Supplementary Figure 5.1 Activation of myelinated sensory neurons by cutaneous and visceral stimuli** Responses recorded in the *Nefh*$^{CreER}$;*R26*$^{LSL\text{-}GCaMP6f}$ mice. Each panel shows deviation projection visualizing cells activated by the stimulus indicated in the row (magenta) and the stimulus indicated in the column (green). Overlap (white) marks neurons activated by both stimuli. The stimulus panel included bladder stretch, colorectal distension (CRD, 80 mmHg), colorectal stroke (CR stroke), anorectal distension (ARD, 80 mmHg), perianal brushing, hair pull, and tail pinch.

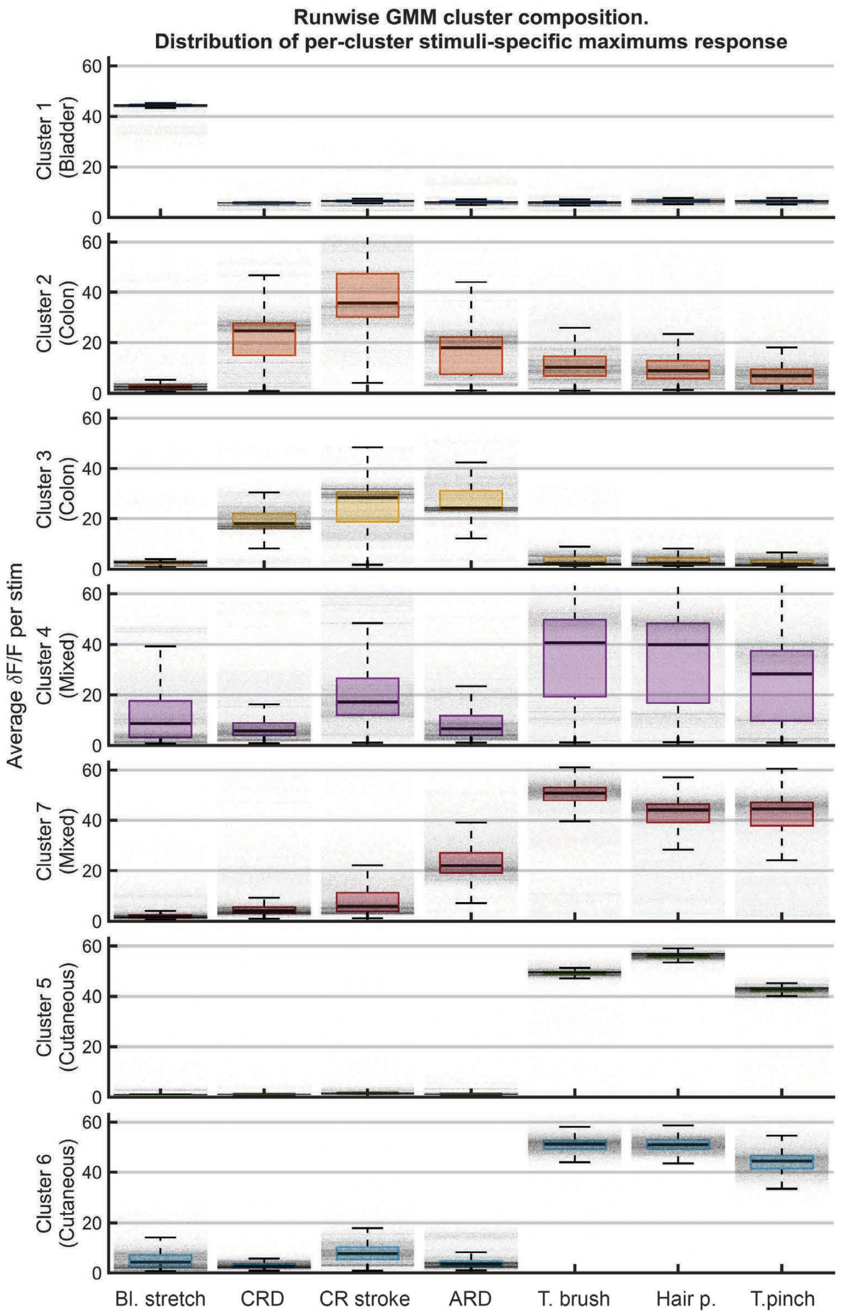


**Supplementary Figure 5.2 Detection of functional cellular modules within myelinated neurons** Stability of GMM cluster composition across 50 000 realizations over data obtained in *Nefh*$^{CreER}$;*R26*$^{LSL\text{-}GCaMP6f}$ model. Box plots show the average stimulus response of cells within each cluster across all realizations; individual dots represent the mean response from a single clustering run. The most representative run was selected for further analysis in Figure 5.

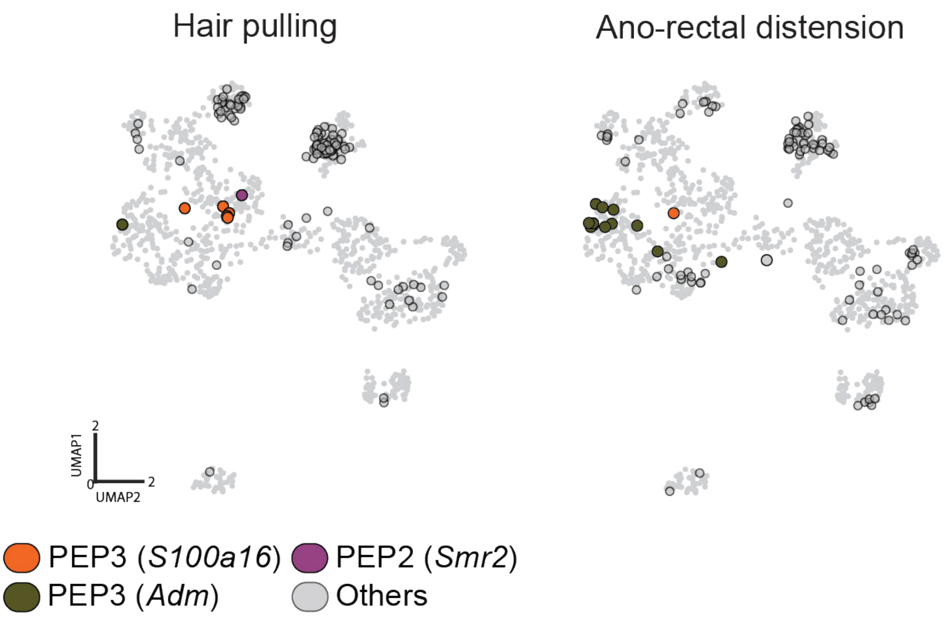


**Supplementary Figure 6.1 A-nociceptors ensemble composition for Hair pulling and Anorectal distension** Overlay of the pelvic DRG transcriptomic atlas (gray) with stimulus-responsive A-nociceptors, color-coded by their class membership.
